# Structural signatures of synergy and redundancy in human brain function

**DOI:** 10.64898/2026.04.09.716459

**Authors:** Laia Barjuan, Maria Pope, M. Ángeles Serrano, Olaf Sporns

**Affiliations:** Departament de Física de la Matèria Condensada, Universitat de Barcelona, 08028 Barcelona, Spain; Universitat de Barcelona Institute of Complex Systems (UBICS), Barcelona, Spain; School of Informatics, Computing & Engineering, Indiana University, Bloomington, IN 47405, USA; Program in Neuroscience, Indiana University, Bloomington, IN 47405, USA; ICREA, Pg. Lluís Companys 23, E-08010 Barcelona, Spain; Department of Psychological & Brain Sciences, Indiana University, Bloomington, IN 47405, USA

## Abstract

A fundamental goal in neuroscience is to understand how the brain’s physical architecture supports complex functional dynamics. While the relationship between structural connectivity and pairwise functional connectivity has been extensively studied, the anatomical basis of higher-order interactions remains poorly understood. In this study, we use multivariate information theory –specifically the O-information– to investigate how the human connectome constrains subsets of brain regions characterized by predominantly redundant or synergistic information sharing. By analyzing the topology and community embedding of these subsets, we reveal two different structural profiles. Redundant subsets are characterized by high internal connection density and strong weights. Their nodes have high clustering and occupy globally less central positions. In contrast, synergistic subsets consist of globally central nodes with high betweenness centrality. We further demonstrate that leveraging these structural features, in particular node centrality, significantly improves the identification of synergistic subsets compared to random sampling. Together, these results demonstrate that the human connectome imposes specific constraints on higher-order information sharing, extending structure–function relationships beyond pairwise interactions and providing new insight into the structural origins of multivariate functional organization.

## 1 Introduction

Unraveling the relationship between structure and function in the brain remains a fundamental challenge in neuroscience, as it addresses how the physical architecture of the nervous system supports complex dynamics and cognition. Structural connections provide the backbone for neural communication and constrain how functional interactions can emerge. Further, this relationship is bidirectional: while structure constrains dynamics, it is also remodeled by long-term functional processes such as learning and plasticity. Yet even in the absence of plasticity, the mapping from structure to function is far from trivial: the same structural scaffold can sustain a rich repertoire of functional patterns that reconfigure over time [1, 2, 3, 4] and, conversely, the same functional pattern may be compatible with a diversity of underlying connectomic configurations through degenerate structure–function mappings [5, 6]. Consequently, considerable effort has been devoted to clarifying the extent to which anatomical connectivity constrains and shapes functional interactions [7, 8, 9, 10, 11, 3, 12, 13, 14, 15, 16, 17].

Network science has provided a powerful framework to investigate this problem by examining the relationship between the network of anatomical connections, termed structural connectivity, and the network of pairwise statistical dependencies between regional activity, termed functional connectivity. Early studies revealed significant correlations between these two networks. For instance, structural and functional connection weights are positively correlated, indicating that regions with strong anatomical links tend to exhibit more synchronous activity [9, 18]. Similarly, nodes that occupy central positions in structural networks are often central in functional networks as well [9]. At a broader scale, it has been observed that dense anatomical connectivity underlies many intrinsic functional systems, especially the visual and somatomotor networks [18, 19].

This approach has provided valuable insights into how structural connectivity influences functional interactions, but it is limited by the inherently pairwise nature with which functional interactions are quantified. Recently, the importance of quantifying higher order functional dependencies in the brain has been emphasized using approaches from topological data analysis [20, 21, 22, 23, 24] and multivariate information theory [25, 26, 27, 28, 29, 30, 31, 32, 33, 34, 35, 36, 37]. In particular, multivariate information-theoretic approaches enable the characterization of higher-order statistical interactions among groups of regions, including the distinction between redundant and synergistic information sharing. Redundant information is duplicated across multiple elements, such that knowledge of one element completely discloses the state of another. In contrast, synergistic information is shared in the joint state of all interacting elements, and is irreducible to any particular element. Recent findings using multivariate information theory show that higher order functional connections in the brain can be both synergistic and redundant, although traditional pairwise functional connectivity analyses have primarily captured redundant relationships [28, 30]. Several studies have highlighted the importance of both modes of information sharing. For instance, the balance between redundancy and synergy has been identified as a predictor of success during auditory discriminatory tasks [32], and characteristic changes in synergy and redundancy have been detected during development and healthy aging [26, 27, 35].

Multiple measures have been proposed to characterize synergy and redundancy, but we are particularly interested in the O-information [38]. This measure is a multivariate information-theoretic quantity whose sign indicates whether interactions among multiple variables are predominantly redundant (when the O-information is positive) or synergistic (when it is negative). The O-information has already been used in studies of the brain [30, 34], and its high scalability makes it a practical and potentially powerful tool for characterizing collective informational structure.

Despite the growing interest in higher-order functional interactions, their anatomical underpinnings, or the extent to which they can be predicted from structural connectivity, remain largely unknown. To date, analyses of this relationship have been limited to the density of structural connections, reporting correlations of both redundancy and synergy with structural connectivity at different anatomical scales [28, 33, 39]. However, a deeper understanding of the anatomical basis of these phenomena, in particular how redundancy and synergy relate to graph-theoretic properties of structural connectivity –such as node centrality or connectivity overlap– rather than to connection density alone, is still missing. Bridging this gap is essential if we want to comprehend not just how function is organized, but how it originates.

In this work, we investigate the structural properties of subsets of brain regions characterized by redundant or synergistic interactions, as quantified by the O-information. We approach this problem from three complementary perspectives: (i) analyzing each subset as an isolated entity, (ii) examining the subset’s embedding within the whole network, and (iii) characterizing the interface between the subset and the rest of the network. Our results show consistent differences between the average structural properties of nodes in subsets that are either synergy or redundancy-dominated. Nodes in redundant subsets tend to be more densely and strongly connected to each other, frequently belonging to the same structural community. On the contrary, synergistic subsets tend to be formed by central nodes–those that are better connected to the rest of the network–, and that are less clearly affiliated with a single community. We also show that leveraging structural information allows us to identify significantly more synergistic subsets than would be expected by chance.

## 2 Results

In our analyses we used data from resting-state fMRI recordings and tractography of two independent datasets: the Human Connectome Project (HCP) [40], comprising 95 participants, and the Microstructure-Informed Connectomics (MICA-MICs) [41], with 50 participants, which served as our replication dataset. In both datasets the cerebral cortex was parcellated into 200 regions using the Schaefer atlas [42]. For each dataset, we concatenated the BOLD time series across participants and runs to enable the computation of information-theoretic measures. The average of the structural connectivity (SC) was constructed using a distance-dependent weighted group-representative [43] procedure. See Methods subsection 4.1 for a detailed description of the acquisition and preprocessing of the datasets and subsection 4.2 for a thorough explanation on the construction of the group-representative networks. The results for HCP will be presented in the main paper and the ones for MICA in the Supplementary Information.

Our aim is to identify characteristic features of structural connectivity in subsets of nodes with synergistic or redundant information-theoretic properties. To this end, we proceed in the following steps. First, we identify subsets of nodes in the network that express high synergy or redundancy. We use the O-information, which extends the notion of mutual information to higher order relationships and captures whether interactions among subsets of nodes are dominated by synergy or redundancy [38]. In particular, the O-information is negative when the dynamics of the subset is dominated by synergistic interactions and it is positive when redundancy prevails. For a full mathematical description of the O-information and the assumptions taken in our calculations see Methods subsections 4.3 and 4.4. Once the relevant subsets have been identified, we use the structural connectivity matrix of the network to characterize their structural features. This approach allows us to determine not only which combinations of nodes display certain higher-order functional properties, but also how these combinations are embedded in the structural network. Thereby, we can link functional higher-order dependencies to the underlying anatomical substrate. An illustrative scheme of our workflow is shown in Fig. 1A.

**Figure 1:**
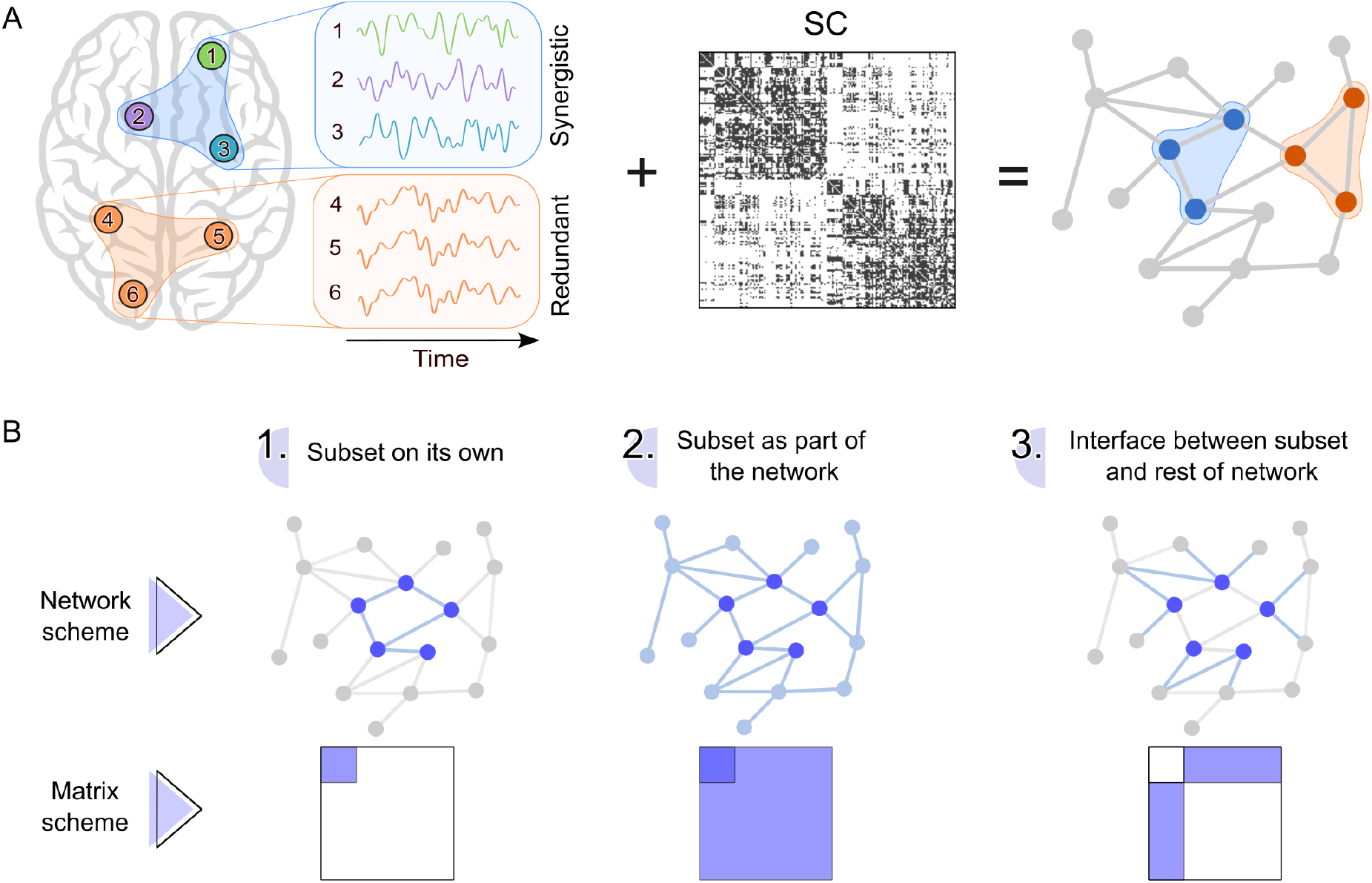
Method and analysis overview. A) Scheme of the approach. From left to right: first, synergistic and redundant subsets of nodes are identified. Then, this information is combined with the structural connectivity matrix (SC) to assess the difference in structural embedding and connectivity patterns of the two types of subset. B) Division of the analysis. From left to right: network components studied at each step. The top row shows the interpretation in terms of nodes and connections. Here, the nodes of the studied subset are highlighted in dark blue, the connections involved are shown in light blue, and disregarded elements are depicted in gray. The bottom row shows the corresponding matrix representation, with selected regions in dark blue and disregarded regions in white.

More specifically, in this study, we compare the structural properties of the most synergistic subsets with those of the most redundant ones, across different subset sizes. Nevertheless, identifying these subsets is computationally hard, as it requires calculating the O-information for all possible subsets –a number that grows combinatorially with subset size. To address this issue, as in previous work [30], we employed a simulated annealing optimization algorithm [44], which allows us to identify both maximally synergistic and maximally redundant subsets in the network, i.e. those with maximally negative and maximally positive O-information, respectively.

The maximally synergistic subsets examined in this paper were previously identified in [30]. In their study, the authors observed that although the O-information of the full network of both datasets is positive (HCP: Ω = 79.16 nats; MICA: Ω = 46.69 nats), it is possible to identify a large number of subsets of nodes with negative O-information for sizes ranging from 3 to 24 (HCP) and from 3 to 27 (MICA). For greater subset sizes, no synergistic subsets were found. On the other hand, it is always possible to identify maximally redundant subsets regardless of the subset size. The maximum redundant subsets used in this paper were not previously published, but they were found using the same methods. Even though in [30] the authors studied subsets with sizes ranging from 3 to 30 nodes, in the current study we analyzed only subsets containing up to 15 nodes. This choice is motivated by the fact that in larger synergistic subsets, nodes belonging to the limbic system –which have a poor signal to noise ratio in fMRI data– tend to dominate and therefore make information measures less reliable. Additionally, some of the analyses are only presented for subsets of size 3, 6 and 10, as they provide a good representation of the entire range.

For each subset size explored, the optimization algorithm was run 5000 times for both synergy and redundancy. Each run is independent and it is possible for multiple runs to converge on the same subset; the frequency of such duplicates reflects the distribution of solutions across the configuration space. For synergy, the number of unique subsets increases rapidly with subset size, from 199 for subsets of size 3 to almost 5000 for subset sizes greater than 13. In particular, the algorithm yields 1582 different results for subset size 6 and 4021 for subset size 10. The specific values can be found in Table ST1 of the Supplementary Information (SI). The picture for redundancy is very different, because the algorithm converges much more reliably onto to the same subsets. In fact, the number of unique subsets for size 3 is 249, which is the maximum value obtained for subset sizes ranging between 3 and 15. For sizes 6 and 10, the values are 185 and 95, respectively. For subsequent analyses, we retained only unique subsets, removing any duplicates. We verified whether matching the number of subsets across all cases affected the results, but similar outcomes were obtained in the vast majority of the analyses. Any exceptions are noted in the text where appropriate. To provide a baseline reference for the comparisons between redundancy and synergy, we generated 5000 randomly selected subsets of nodes for each subset size.

One could argue that using an optimization algorithm to find the maximally synergistic and maximally redundant subsets is unnecessary. To justify its use, we generated 5 million subsets of nodes at random, for each subset size. We calculated the O-information value for all of them, kept the 1000 subsets in both extremes of the O-information spectrum as our synergistic and redundant subsets and ran the analyses for these subsets. This approach was able to approximately reproduce the observed results for synergistic subsets, but it failed at replicating the ones for redundant subsets. Only for subset size 3, the results were comparable. In addition, the subsets obtained through this method exhibited absolute O-information values smaller than those identified by the optimization algorithm, for both synergistic and redundant cases, though the difference was particularly pronounced for the redundant ones. Therefore, random sampling is unable to identify, especially, the maximally redundant subsets present in the network, making the optimization algorithm essential for our purposes.

### 2.1 Mean structural properties

We approached the analysis of the structural properties of the functional subsets in three ways. First, we focused on the subset as a separate entity, disregarding its connections to the rest of the network. Second, we studied the subset taking into account its embedding in the rest of the network. And finally, we examined the interface between the subset and the rest of the network (i.e., the subset’s inputs and outputs). In Fig. 1B, we provide a graphical, qualitative illustration of what each approach represents in terms of the nodes and connections of the network involved. Additionally, we present a schematic representation of the corresponding areas in the structural connectivity matrix.

In all cases, we examined the distributions of the mean structural properties associated with each subset, distinguishing between redundant, synergistic, and randomly selected subsets. The statistical significance of the differences between each pair of distributions –synergistic–redundant, synergistic–random, and redundant–random– for subset sizes 3, 6 and 10 was assessed using a combination of the Kolmogorov–Smirnov (KS) test and a permutation test. In particular, we calculated the p-values by comparing the KS statistic of the original distributions (D) and those of 10^4^ instances of a null model obtained by randomly shuffling the data labels. For discrete distributions, instead of using the KS statistic for the comparison, we used the mean. The p-values are reported throughout the text and the specific values of the experimental KS statistics are reported in the SI.

#### 2.1.1 Subset on its own

We first studied the subsets independently to the rest of the network. Our goal was to determine whether the subsets exhibit specific connectional properties that reflect the capacity for information exchange among their constituent nodes. In particular, we examined the number of connected components within the subset, its connection density and its connection weights.

Redundant subsets form one connected component more frequently than synergistic subsets, and synergistic subsets form one connected component more frequently than randomly selected subsets of nodes up to approximately subset size 12, Figs. 2A and 2D. These results are directly related to the density of connections, Figs. 2B and 2E. Here we can see that, for all subset sizes, redundant subsets have a much higher density of connections as compared to random subsets. Meanwhile, the connection density of synergistic subsets decreases with subset size, with mean values remaining higher than those of random subsets up to approximately size 10. In Fig. 2E, we see that despite having the same median value of connection density for subset size 3, the distributions of synergistic and redundant subsets gradually drift apart as subset size increases. Additionally, nodes in redundant subsets are connected by stronger connections than nodes in synergistic subsets, which, in turn, are connected by stronger connections than expected at random. This behavior can be seen in Fig. 2C and, especially, in Fig. 2F, where the distinction between synergy and redundancy becomes increasingly evident as the number of nodes in the subset grows. All p-values corresponding to the comparisons between synergistic–redundant, synergistic–random, and redundant–random distributions, for the subset sizes shown in the figure are *p* = 0. The results obtained using the MICA dataset (Fig. SF2 in the SI) are consistent with those shown in this figure (all *p <* 6 · 10^−4^).

**Figure 2:**
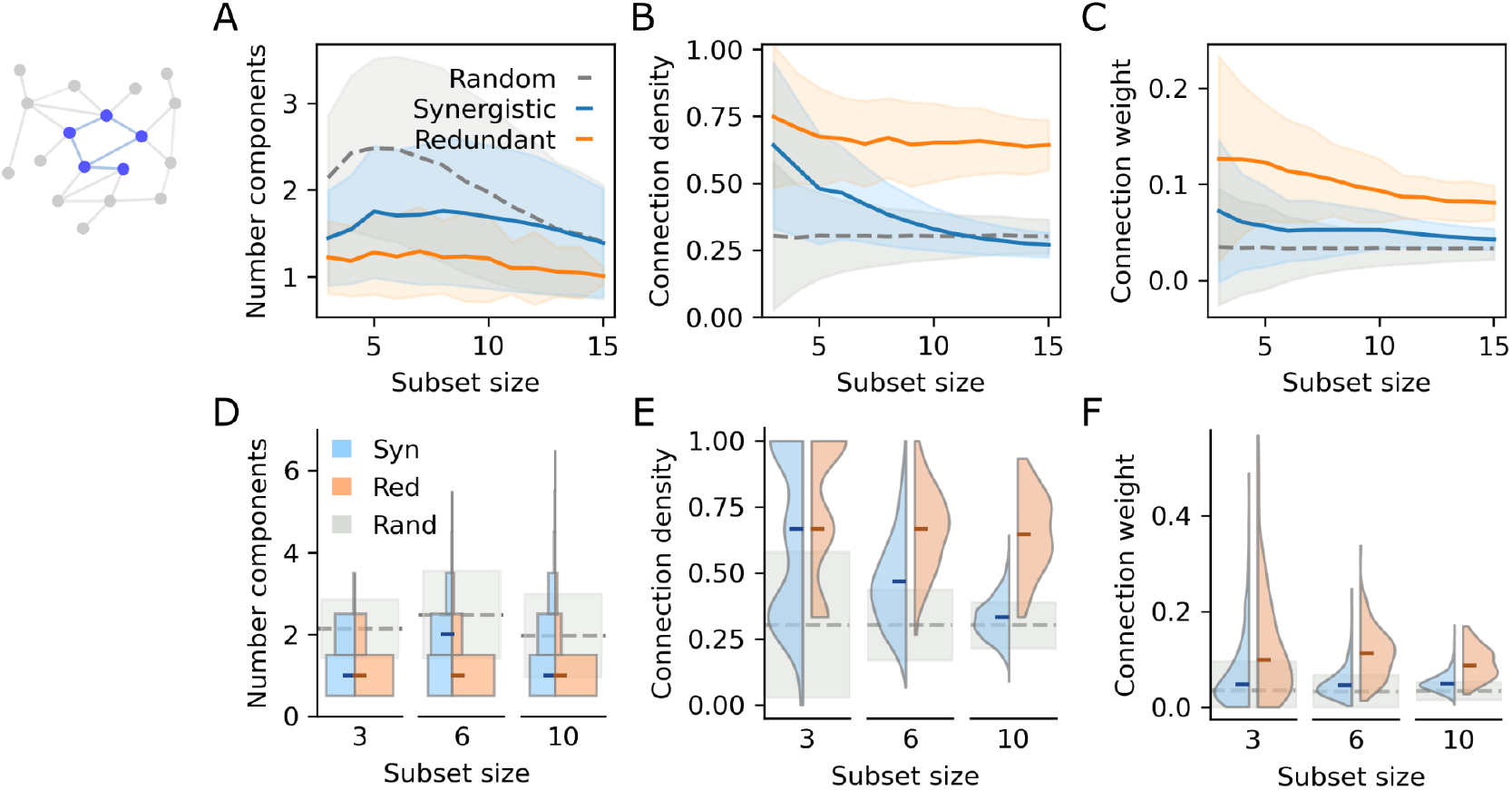
Subset on its own. A-C) Mean results of A) number of components within the subgraph, B) connection density, and C) average connection weight, across all subset sizes. The results for synergistic subsets are represented by the solid blue line; for redundant subsets by the solid orange line; and for random subsets by the dashed gray line. The shadowed areas correspond to the standard deviation in each case. D-F) Distributions of values of D) number of components, E) connection density and F) average connection weight, for subsets with 3, 6 and 10 nodes. Values for synergistic and redundant subsets are depicted in blue and orange, respectively. The median value of each distribution is indicated by a small horizontal mark. The mean and standard deviation for random subsets are represented by the dashed gray line and the shadowed gray area, respectively.

In brief, redundancy-dominated subsets are characterized by densely and strongly interconnected architectures that support direct signal transmission. Although synergy-dominated subsets are less tightly connected, their topology enables more efficient communication than in randomly sampled subsets.

#### 2.1.2 Subset as part of the network

Next, we studied the properties of the subsets in the context of their embedding within the whole brain network. This analysis is based on the assumption that a subset’s informational properties depend not only on the subset itself, but also on how its nodes are positioned within the broader network, i.e. how they connect to or remain isolated from the rest of the system. To address this, we examined whether nodes in each type of subset differ in terms of centrality and cohesion. In particular, we calculated the average degree, strength, betweenness centrality, and clustering coefficient of the nodes constituting each subset.

Nodes in synergistic subsets have higher average degree than nodes in redundant subsets, while random subsets lie in between. Above subset size 10, however, the average for synergistic subsets falls below that of random subsets too, see Figs. 3A and 3E. Interestingly, as subset size increases, the separation between synergistic and redundant distributions for HCP becomes less pronounced, although the overall trend persists (all p-values equal to 0). For MICA, in contrast, the distributions remain distinct as subset size grows, Fig. SF3 of the SI. The distributions of the average node strength in synergistic and redundant subsets overlap substantially at small subset sizes –both having a slightly higher average strength than their random counterparts– and become somewhat more distinct as more nodes are considered, see Figs 3B and 3F (all p-values less than 7 *·* 10^−4^). The distinction at larger subset sizes is particularly pronounced for MICA, while it is much less evident in HCP. In Figs. 3C and 3G, we see that the average betweenness centrality of nodes in synergistic subsets is higher than in redundant subsets, with random subsets lying in between (all p-values less than 7 · 10^−4^). As observed for degree, beginning at subset size 10, the values for synergistic subsets drop briefly below those of random subsets, yet remain higher than those of redundant subsets. Finally, the average clustering coefficient of nodes in synergistic subsets is lower than in redundant subsets, with random subsets falling again in an intermediate range, see Figs. 3D and 3H (all *p* = 0). The results obtained using the MICA dataset (Fig. SF3 in the SI) are consistent with those shown in this figure, except where already noted.

**Figure 3:**
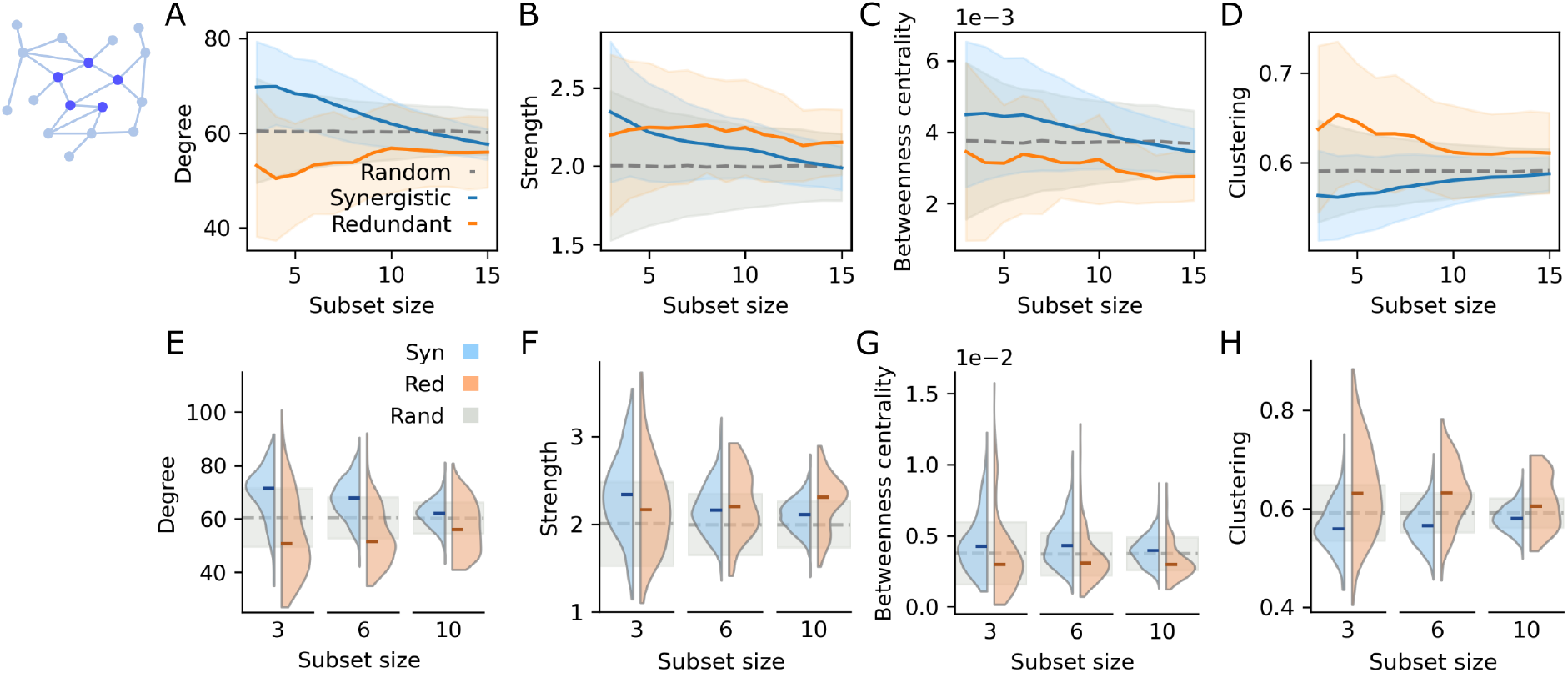
Subset as part of the network. A-D) Mean results of average A) degree, B) strength, C) betweenness centrality and D) clustering coefficient, across all subset sizes. The results for synergistic subsets are represented by the solid blue line; for redundant subsets by the solid orange line; and for random subsets by the dashed gray line. The shadowed areas correspond to the standard deviation in each case. E-H) Distributions of average E) degree, F) strength, G) betweenness centrality and H) clustering coefficient, for subsets with 3, 6 and 10 nodes. Values for synergistic and redundant subsets are depicted in blue and orange, respectively. The median value of each distribution is indicated by a small horizontal mark. The mean and standard deviation for nodes in random subsets are represented by the dashed gray line and the shadowed gray area, respectively.

In summary, redundant subsets consist of nodes that are globally less central but locally integrated, while synergistic subsets comprise nodes with enhanced global connectivity and reduced local integration.

In previous work [30], subsets comprising nodes from 6 or 7 distinct functional communities were more likely to be synergistic than subsets drawn from fewer communities. Here, we tested whether a similar pattern emerges when considering structural communities identified with multiresolution consensus clustering (MRCC), see Methods section 4.5 for more details. This approach yielded 10 structural communities of varying sizes, which can be seen in the SC matrix ordered by communities displayed in Fig. 4A. We found that, on average, redundant subsets draw from a smaller number of structural communities compared to synergistic subsets, especially as subset size increases, see Figs. 4B and 4E (all p-values equal to 0).

**Figure 4:**
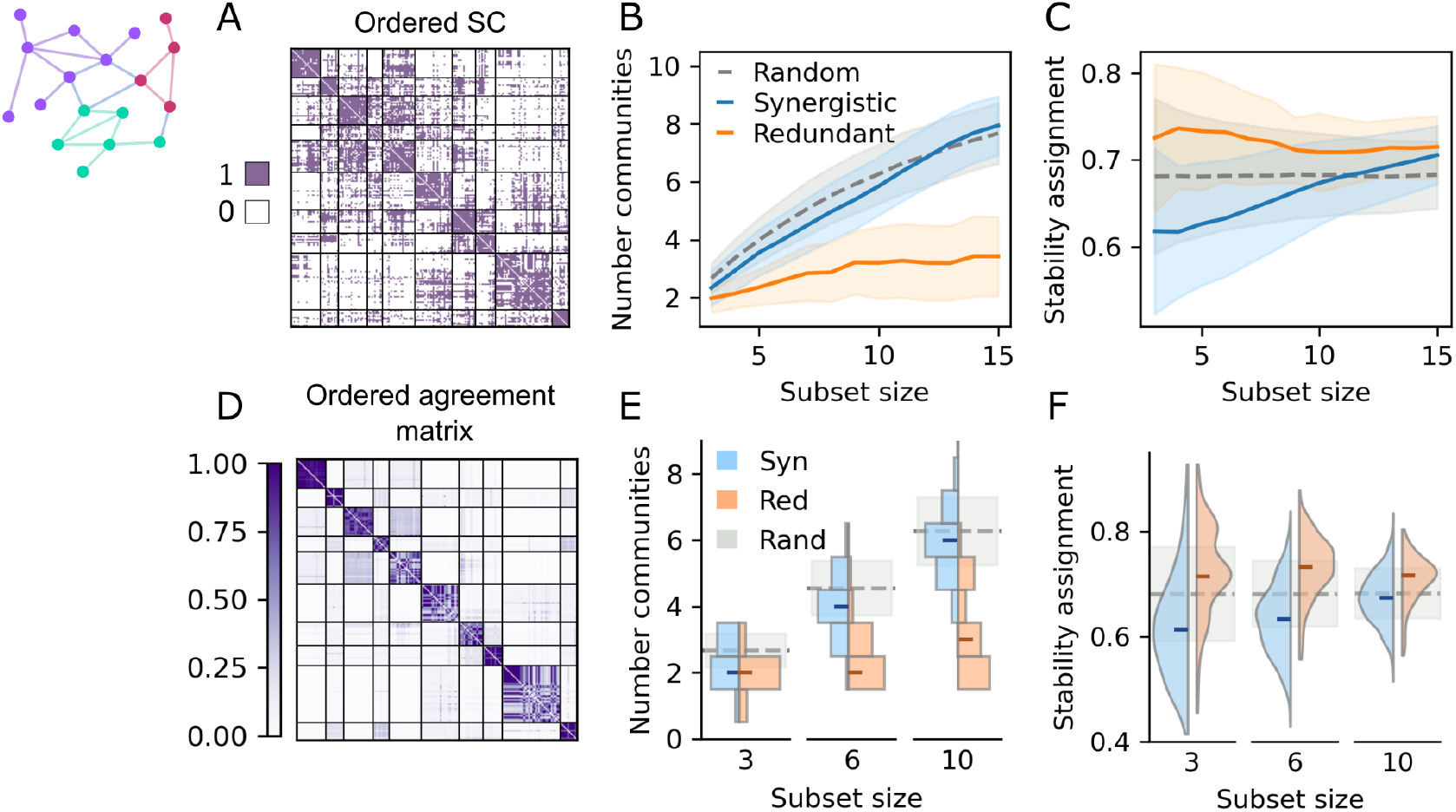
Community structure. A) Structural connectivity matrix ordered by structural communities, whose boundaries are indicated by black solid lines. Existing connections are shown in purple; non-existing connections in white. B-C) Mean results of number of communities appearing in each type of subset and of node community assignment stability, respectively. The results for synergistic subsets are represented by the solid blue line; for redundant subsets by the solid orange line; and for random subsets by the dashed gray line. The shadowed areas correspond to the standard deviation in each case. D) Agreement matrix ordered by structural communities, whose boundaries are indicated by black solid lines. Darker colors represent higher agreement. For subsets with 3, 6 and 10 nodes: distributions of E) number of communities appearing in each type of subset and F) average node community assignment stability. Results for synergy and redundancy are shown in blue and orange, respectively. The mean and standard deviation for random subsets are represented by the dashed gray line and the shadowed gray area, respectively.

As part of the consensus procedure, we get an agreement matrix –whose entries reflect how often each pair of nodes is assigned to the same community across spatial resolutions–, which we display in Fig. 4D. From the agreement matrix, we calculated the node-level stability of community assignment as the average agreement between a given node and the other nodes within its community. In Figs. 4C and 4F we can see that nodes in redundant subsets are assigned more reliably to the same community than nodes in synergistic subsets, with their mean values lying above and below the average of the network, respectively (all p-values equal to 0). Notably, beginning at subset size 10, the average for synergistic subsets starts to exceed the network average too. These results suggest that, particularly for small to moderate subset sizes, nodes involved in redundancy have a clear affiliation with a particular module, while nodes participating in synergy plausibly fit in multiple structural communities.

The results for MICA are shown in Fig. SF4 of the SI and follow the same general trend as for HCP.

#### 2.1.3 Interface between subset and rest of the network

In the previous section, we assessed the positioning of the nodes in each subset within the broader network from a centrality, cohesiveness and community membership perspective. To gain a complete understanding of how subsets are connected to, or isolated from, the rest of the system, we next examine the interface between each subset and the remaining network. Specifically, we study the subset-to-network connection density, the fraction of nodes of the network that are one connection away from the nodes in the subset, and the average similarity between the neighbors of the nodes in the subset (excluding those neighbors within the subset). We quantified this neighbor similarity using the Jaccard index between connection patterns, defined as the number of shared neighbors divided by the total number of unique neighbors across each node pair.

The subset-to-network connection density is the number of links between the nodes in the subset and the other nodes of the network divided by the total number of possible links. Synergistic subsets are more densely connected to the rest of the network than redundant subsets, while random subsets lay in between, see Figs. 5A and 5D (all *p* = 0). However, the difference in the median value of the distributions decreases as the size of the subset grows, with the average for synergistic subsets falling below the network average at subset size 10.

**Figure 5:**
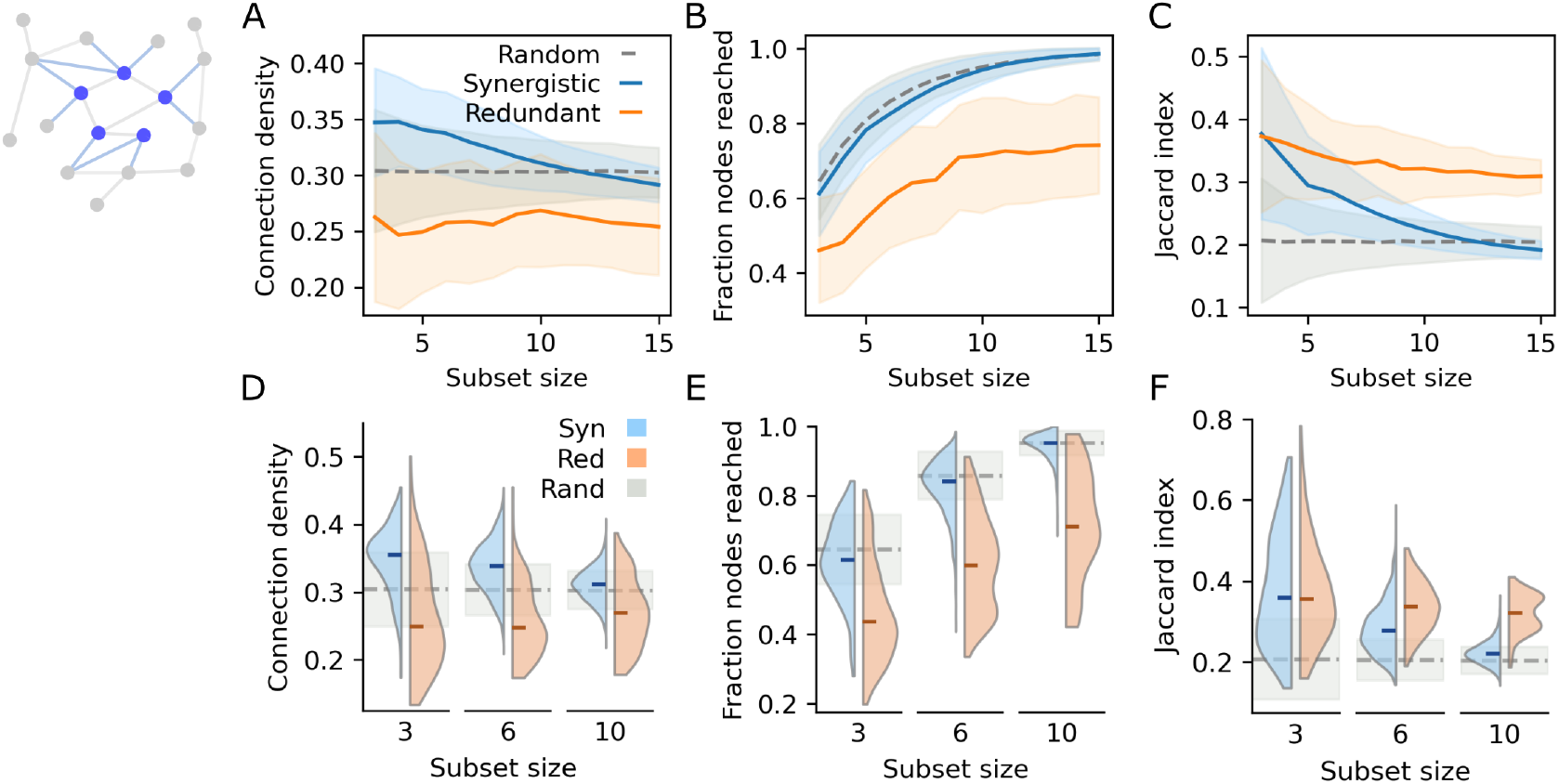
Interface between subset and rest of network. A-C) Mean results of A) subset-to-network connection density in the interface, B) fraction of nodes of the network that are one connection away from the nodes in the subset, and C) average Jaccard index between neighbor patterns. The results for synergistic subsets are represented by the solid blue line; for redundant subsets by the solid orange line; and for random subsets by the dashed gray line. The shadowed areas correspond to the standard deviation in each case. For subsets with 3, 6 and 10 nodes: distributions of D) subset-to-network connection density in the interface, E) fraction of nodes of the network that are one connection away from the nodes in the subset, and F) average Jaccard index between neighbor patterns. Results for synergistic and redundant subsets are shown in blue and orange, respectively. The median value of each distribution is indicated by a small horizontal mark. For random subsets, the mean is indicated by the dashed gray line and the standard deviation by the shadowed gray area.

We also analyzed the fraction of nodes of the network that are one connection away from the nodes in the subset. Consistently across subset sizes, synergistic subsets reach a significantly larger fraction of nodes than redundant subsets, see Figs. 5B and 5E (all *p <* 4 · 10^−4^). We also observe that while the variability of number of nodes reached decreases with size for synergistic subsets, it remains large for redundant subsets.

Finally, we computed the average Jaccard index between the rows of the SC matrix corresponding to the nodes in the subset, excluding connections within the subset itself. The portion of the matrix used is illustrated in column 3 of Fig. 1B. Conceptually, this corresponds to measuring the number of shared neighbors divided by the total number of unique neighbors for each pair of nodes in the subset, and then averaging across all pairs. Thus, this metric quantifies the average similarity in interface neighbors among nodes in the subset. We observe that the distributions of the average Jaccard index for synergistic and redundant subsets are very broad and overlap substantially for small subset sizes. In fact, at subset size 3, they are not significantly different (*p* = 0.1219), i.e. synergistic and redundant subsets follow the same distribution. All other p-values are equal to 0. The median value of both synergistic and redundant distributions is higher than expected at random for small subset sizes. However, as subset size increases, the distributions gradually separate, with the one for synergistic subsets moving closer to the random case, see Figs. 5C and 5F.

To summarize, redundant subsets are relatively less connected to the rest of the network and tend to interface repeatedly with the same external nodes. In contrast, synergistic subsets are broadly connected, establishing links with nearly all parts of the network.

The results for MICA are shown in Fig. SF5 of the SI and are consistent with those presented here (with all *p <* 10^−2^).

### 2.2 Participation in subsets

To better characterize which elements of the network contribute to each type of subset, we quantified the percentage of nodes and connections that appear at least once in the optimized synergistic and optimized redundant subsets. A connection was considered part of a subset if it linked two nodes included in that subset. Importantly, this definition does not imply that the connection itself directly mediates the communication dynamics that give rise to synergy or redundancy; it only reflects the existence of the physical connection within the subset. It is also important to note that these results are influenced by the number of unique subsets considered for each subset type and size. This variability arises from both the distribution of solutions across the configuration space and the convergence of the optimization algorithm, as mentioned at the beginning of the Results section. Figure 6A shows that the percentage of nodes participating in optimized redundant subsets is smaller than in optimized synergistic subsets, where almost the entirety of nodes appear at least once for subset sizes greater than 5. Likewise, fewer edges of the network appear at least once in optimized redundant subsets, see Fig. 6B. Specifically, the number of connections comprised in redundant subsets stays fairly constant around 10% across subset sizes, while in synergistic subsets it increases from approximately 10% when subsets contain 3 nodes to around 50% when they contain 15. In Fig. 6C we show, for subset size 6, which connections appear within subsets (within only synergistic or redundant subsets, or both), as well as connections that do not appear in either. It is worth noting that the connections appearing in synergy and in redundancy overlap only very infrequently. In fact, as we can observe in the bottom-left matrix, very few connections appear both in synergistic and redundant subsets. At the same time, most of the connections are never involved in any kind of subset for this subset size, bottom right. Results from the MICA dataset (Fig. SF6, SI) closely match those presented in this figure.

**Figure 6:**
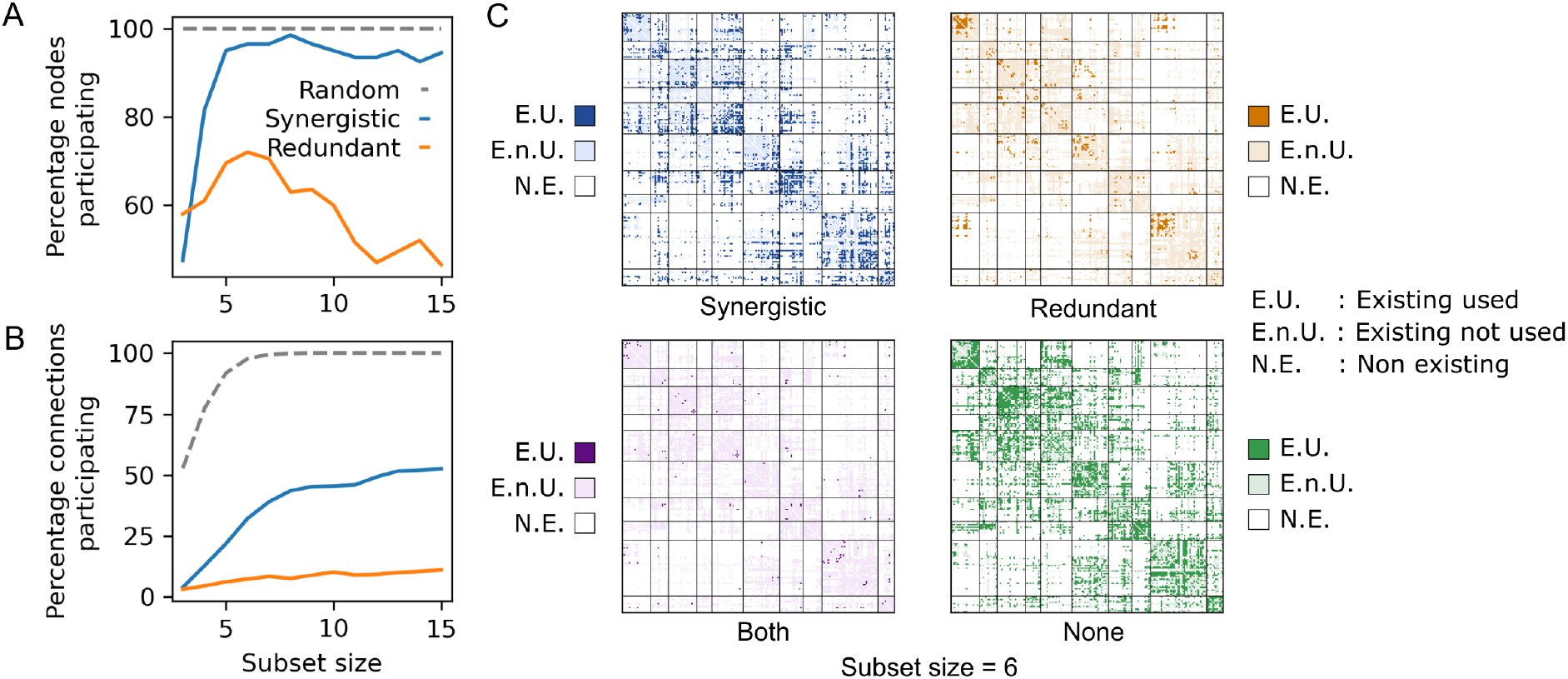
Participation of nodes and connections in optimized subsets. Percentage of A) nodes and B) connections participating at least once in each type of subset across all subset sizes. Results for synergistic, redundant and random subsets are shown by the solid blue line, solid orange line and dashed gray line, respectively. C) Instances of the SC matrix where connections that meet a certain criteria are highlighted: in the top-left panel, connections participating in synergistic subsets; in the top-right, those in redundant subsets; in the bottom-left, those in both; and in the bottom-right, those that never participate. In all panels, non-existent connections are shown in white, existing but non-highlighted connections in light colors, and highlighted connections in bright colors. Results correspond to subsets with 6 nodes.

We emphasize once more that, despite the results presented above, it would be inaccurate to claim that fewer nodes or connections participate in redundant subsets as compared to synergistic subsets. A more precise statement is that the optimized redundant subsets involve fewer nodes and connections than the optimized synergistic subsets. In other words, maximal redundancy is concentrated within more specific subsets of nodes, whereas maximal synergy is distributed more heterogeneously across the network. The equivalent results calculated matching the number of subsets can be found in Fig. SF1 of the SI.

In previous sections, we have compared the average structural properties of the nodes in synergistic, redundant, and random subsets at the group level. However, a previous study [30] found that brain regions do not equally participate in synergistic and redundant subsets. To determine whether this unequal involvement is related to network structure, we next examined participation at the node level: we quantified how often each node appeared in each subset type and tested whether this participation frequency correlated with its structural properties. To quantify the correlation we used the Spearman coefficient. In particular, we computed the correlation between participation frequency and degree, strength, betweenness centrality (BC), weighted betweenness centrality, clustering, weighted clustering and the stability in the community assignment. The results are shown in Fig. 7A. For synergistic subsets, we observe a positive correlation between participation frequency and some of the nodes’ structural properties, like the degree and the betweenness centrality. The correlation with their weighted counterparts, i.e. strength and weighted betweenness centrality, is less pronounced. Moreover, we also obtain an inverse correlation between participation frequency and the stability in the community assignment, and less prominently between participation frequency and clustering. Interestingly, all these correlations decrease steadily with subset size. For redundant subsets, on the other hand, we only find a weak correlation between participation frequency and the weighted measures, i.e. strength, weighted betweenness centrality and weighted clustering. This suggests a clear association between the strongest connections in the network and redundancy, as we also observed in Figs. 2C and 2F.

**Figure 7:**
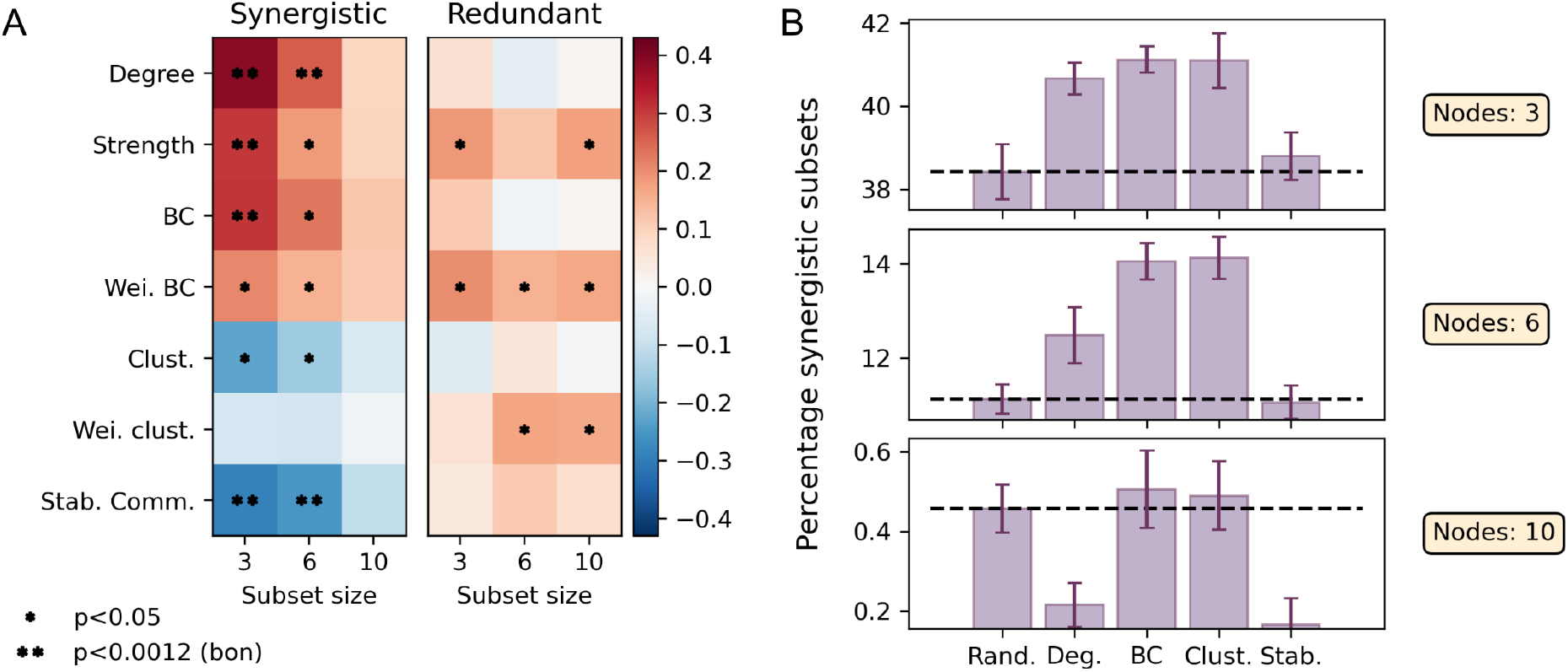
Node participation frequency. A) For synergistic and redundant subsets with 3, 6 and 10 nodes: Spearman correlation between node participation frequency and degree, strength, betweenness centrality (BC), weighted betweenness centrality, clustering, weighted clustering and stability in community assignment. Positive and negative correlations are shown in red and blue, respectively. Entries with p-values smaller than 0.05 are marked with an asterisk. Entries with p-values smaller than the Bonferroni-corrected threshold (*p* = 0.0012) are marked with two asterisks. B) Percentage of synergistic subsets obtained biasing the selection process by degree (Deg.), betweenness centrality (BC), clustering (Clust.) and stability in community assignment (Stab.) with respect to random selection. The limits of the y-axis have been adjusted to better visualize the differences between bars.

#### 2.2.1 Biased selection of synergistic subsets

The results presented above show that a node’s structural properties strongly predict whether it preferentially participates in synergistic subsets. Is it then possible to use those properties to build synergistic subsets *de novo*? To assess this hypothesis, we performed a biased selection of subsets based on the unweighted structural properties of the nodes. More specifically, we generated subsets by selecting nodes from the top (bottom) two quartiles of magnitudes that were positively (negatively) correlated with participation frequency. Then, we calculated the percentage of synergistic subsets that we obtained in each case and we compared our results to pure random selection. To reduce the effect of stochasticity in the subsets’ construction, the results were averaged over 10 independent realizations of a 5000-subset selection. In Fig. 7B, we show the percentage of synergistic subsets obtained at random and with the biasing method, for subsets with size 3, 6 and 10. We observe that, especially for small subset sizes, the biasing approach increases our chances of identifying subsets with synergistic properties. In particular, the betweenness centrality and clustering seem to give the best performance, regardless of subset size. This effect is less prominent for big subset sizes, and disappears for subsets with more than 10 nodes.

The corresponding results for the MICA dataset are shown in Fig. SF7 of the SI. In general, the results are consistent. We obtain patterns for the synergistic subsets similar to those in Fig. 7A, although the correlations are less significant than for HCP. In particular, the negative correlation between participation in synergy and community stability fades. For redundancy, the positive correlation between participation frequency and the weighted measures remains, although less significant, but in this case we also observe a clear correlation between redundancy and community stability. In the MICA dataset, the biasing of the selection process does not increase the percentage of synergistic subsets as strongly as in HCP, although betweenness centrality (BC) still yields a modest improvement. However, when restricting our analysis to the strongly synergistic subsets–those with O-information values sufficiently below zero (*τ* = −0.01)– the improvement becomes more apparent.

## 3 Discussion

In this paper, we trace patterns of multivariate information in the human brain, such as synergy and redundancy, to specific features of the underlying structural connectivity. We show that synergistic and redundant subsets are rooted in distinct anatomical architectures: nodes participating in redundant subsets are locally integrated but globally peripheral, while those in synergistic subsets act as globally connected bridges across structural communities.

Focusing first on redundancy, we find that optimized redundant subsets are dense, strongly connected units that usually form a single connected component with above-average internal connection weight. This suggests that maximal redundancy relies on tight wiring for efficient direct communication. Additionally, redundant subsets consist of nodes with lower degree and betweenness centrality, yet higher clustering than the network average. They are poorly connected to the rest of the network, and their nodes often share neighbors. Therefore, redundant subsets are locally well-integrated subnetworks that may act as information reservoirs, built for stable, perturbation-resistant internal communication, rather than network-wide information spreading. Moreover, each subset draws nodes from few structural communities, and its nodes show high consistency in their community assignments, indicating that redundancy concentrates in structurally cohesive modules.

Like redundant subsets, optimized synergistic subsets are internally dense and have slightly above-average connection weights. Moreover, they form a single connected component more often than expected by chance, suggesting that they are also topologically suited for efficient communication and within-subset integration. In contrast, they are more heterogeneous than redundant subsets, drawing nodes from a broader range of structural communities and that are, individually, less cleanly assigned to any single community. This points to synergy emerging from structurally diverse nodes that can integrate information across communities. Consistent with this interpretation, synergistic subsets are densely connected to the rest of the network and are composed of nodes with high degree, high betweenness centrality, and low clustering, i.e., nodes that are globally well-connected and strategically positioned to combine distinct informational streams.

Some findings align with previous observations by Luppi et al. [28], who reported that redundancy is more closely linked to structural connectivity than synergy, with similarly wired regions sharing redundant information and regions with diverse profiles being more prone to synergy. However, they identified interactions using Integrated Information Decomposition, which quantifies dependencies between the past and future of co-evolving pairs of regions. Consequently, theirs is a measure of temporal redundancy and synergy, more closely related to emergence, and limited to pairwise interactions across brain regions [45]. It is currently unknown if or how results from temporal synergy and redundancy relate to those without a temporal element. By contrast, our approach characterizes spatially multivariate dependencies, allowing us to investigate redundant and synergistic information shared across larger sets of regions.

While our initial analysis focused on subset averages, previous work showed that some nodes participate frequently in a given type of subset, whereas others appear only occasionally [30]. This variability raises the question of whether participation frequency reflects specific structural properties. In synergistic subsets, node participation frequency correlates positively with centrality metrics (degree, strength, and betweenness centrality) and negatively with cohesion-related attributes (clustering and stability in community assignment). This indicates that as the node’s centrality increases, so does its probability of participating in synergistic processes. In redundant subsets, node participation frequency correlates positively with weighted structural properties (strength, weighted betweenness centrality, and weighted clustering), highlighting the key role of connection weight in redundant interactions.

Synergistic subsets are considerably less frequent than redundant ones [30], making their identification via random sampling very unlikely. Consequently, the structural signatures we identified in synergistic subsets are particularly important because they can guide subset selection. Specifically, combining nodes with high betweenness centrality –those most involved in efficient communication paths– increases the probability of detecting synergistic subsets. In other words, the combination of multiple nodes strategically positioned to mediate cross-module flows facilitates synergy, which is consistent with the idea that synergy reflects integration of information across distributed sources. These results further highlight the practical importance of structural factors in predicting information properties. However, not all subsets with the appropriate structural features exhibit synergy, suggesting that additional, yet-to-be-explored factors contribute to the emergence of synergistic behavior.

A general observation from our results is that the average structural properties of nodes forming redundant subsets remain stable across subset sizes, whereas nodes in synergistic subsets exhibit increasing randomness as subset size grows. When approximately ten nodes are involved, the structural properties of nodes in synergistic interactions resemble those expected by chance, indicating a loss of structural specificity. This contrast can be traced back to the composition of the subsets: nodes in redundant subsets are consistently reused across subset sizes, while nodes in synergistic subsets are gradually replaced as the size of the subset grows. This suggests that large-scale synergistic interactions may rely less on specific structural constraints or, as discussed in [34], may be increasingly dominated by noise.

Taken together, our findings show that higher-order interactions are not distributed randomly across the brain but are rooted in a specific anatomical basis. Redundancy is primarily a local phenomenon, thriving in “short-path” neighborhoods, consistent with previous findings about it being captured by pairwise statistics [28, 30]. Synergy, however, involves nodes that occupy central positions in the system’s global communication, supporting interactions that cannot be reduced to simple pairwise dependencies.

It has been established that higher-order structures support higher-order functions [46], but are explicitly higher-order mechanisms necessary for observing higher-order statistics? Our results suggest that particular patterns of pairwise connectivity may be sufficient to generate higher-order information dynamics. While we cannot rule out the possibility that higher-order mechanisms contribute to the observed information-theoretic patterns, no such mechanisms are currently known to operate at this scale of brain organization. Our findings thus complement theoretical efforts aimed at disentangling the origins of higher-order information structure [47].

One limitation of our study is that estimating O-information reliably requires using group-averaged data, which removes individual differences and may generate structural connectivity patterns that do not match any single subject, potentially biasing inferred structural features. In addition, O-information measures the dominance of synergy or redundancy, rather than their exclusivity. Consequently, labeling a subset as redundant does not imply the absence of synergistic information within it. In fact, in [34] it was found that the same subsets experience both the most strongly synergistic and strongly redundant time points. Since we treated subsets as strictly dominated by one regime, we cannot tell how the wiring of a synergy-dominated subset supports any redundancy it contains, or vice versa, an issue that future information decompositions on small subsets could address [48, 49, 50, 45]. Finally, we measured anatomical edges coincident on nodes contained within subsets. However, the existence of an edge within a subset does not necessarily imply its participation in functional interactions. This distinction cannot be resolved with current neuroimaging techniques, but our analysis remains valuable as it reveals consistent anatomical connectivity differences between synergistic and redundant subsets.

Future work could explore the overlap between synergistic and redundant subsets at both the connection and node levels. Identifying regions that frequently lie at these intersections could provide insights into their functional roles.

In conclusion, specific structural features of the human connectome predispose nodes to participate in synergistic or redundant interactions. Redundant subsets form tightly connected configurations that are ideal for reliable information maintenance, whereas synergistic subsets comprise nodes ideally positioned to integrate information across the network. These complementary architectures highlight the key role of structural connectivity in shaping the brain’s higher-order information dynamics.

## 4 Methods

### 4.1 Description of the datasets

This study was performed with a dataset derived from the Human Connectome Project [40] (HCP), and the results were replicated using the MICA-MICs dataset [41] (MICA). Both datasets were parcellated into 200 cortical regions (nodes) using the Schaefer atlas [42].

#### Human Connectome Project

The original HCP dataset consisted of 100 unrelated subjects, but 5 were excluded due to motion criteria developed before the present study. Subjects that exceeded 1.5 times the interquartile range of the distribution of mean and mean absolute deviation of the relative root mean square of motion were excluded (a total of four subjects) and a fifth was excluded due to a software error during diffusion MRI processing. All participants provided informed consent to data procedure protocols approved by the Washington University Institutional Review Board. The remaining subset of 95 subjects were formed by 56% of female participants and the mean age of the cohort was 29 *±*4.

A thorough description of HCP dataset data acquisition and preprocessing can be found in [40, 51]. As a summary, the data collection for the HCP were conducted on a Siemens 3T Connectom Skyra equipped with a 32-channel head coil. Structural connectivity (SC) was derived from diffusion imaging and tractography, using two diffusion MRI scans per subject acquired with a spin-echo planar imaging sequence (repetition time [TR] = 5520 ms, echo time [TE] = 89.5 ms, flip angle = 78°, 1.25 mm isotropic voxel resolution, b-values = 1000, 2000, 3000 s/mm2, 90 diffusion weighed volumes for each shell, 18 b = 0 volumes). The two scans were taken with opposite phase encoding directions and averaged. Functional Connectivity (FC) was derived from resting-state functional MRI (rs-fMRI), acquired with a gradient-echo echo-planar imaging (EPI) (repetition [TR] = 720 ms, echo time [TE] = 33.1 ms, flip angle = 52°, 2 mm isotropic voxel resolution, and multiband factor of 8). In total, four scans were taken on two separate days, and each scan was 14:33 min long. Participants were asked to keep their eyes open.

The preprocessing of the diffusion images included normalization to the mean b0 image, correction for EPI, eddy current, and gradient nonlinearity distortions, correction for motion, alignment to the subject anatomical space using a boundary-based registration [52], and intensity nonuniformity correction. For each subject, multiple runs of probabilistic tractography were conducted using Dipy’s Local Tracking module, with turning angle fixed at 20° for step sizes 0.25 mm, 0.4 mm, 0.5 mm, 0.6 mm, and 0.75 mm, and with step size fixed at 0.5 mm for turning angles of 10°, 16°, 24°, and 30°. Within each voxel of a white matter mask, streamlines were randomly seeded three times, and they were deleted if they were shorter than 10 mm or terminated in anatomically implausible locations. Erroneous streamlines were subsequently filtered based on the cluster confidence index [53]. The number of streamlines connecting different regions in the parcellation was recorded for each tractography run, and it was normalized by the geometric mean of the volumes of the connected nodes. The values of the edges were calculated as the weighted mean across the 9 runs of the tractography algorithm, where weights reflected the proportion of total streamlines contributing to each edge. This normalized number of streamlines is what we refer as connection weight.

The preprocessing of rs-fMRI data included [51] distortion, susceptibility, and motion correction, registration to subjects’ respective T1-weighted data, bias and intensity normalization, projection onto the 32k fs LR mesh and alignment to common space with a multimodal surface registration [54]. This preprocessing resulted in ICA+FIX time series in the CIFTI grayordinate coordinate system. Further preprocessing included global signal regression and detrending and band pass filtering (0.008 to 0.08 Hz) [55]. The first and last 50 frames were discarded after confound regression and filtering, leaving 1100 frames corresponding to a total scan duration of 13.2 minutes.

#### MICA-MICs

The MICA dataset consists of 50 unrelated subjects. All participants provided informed consent and the procedures and protocols were approved by the Ethics Committee of the Montreal Neurological Institute and Hospital. The participants were formed by 46% of female participants and the mean age of the cohort was 30 ±6.

A thorough description of MICA dataset data acquisition and preprocessing can be found in [41]. As a summary, the data collection for the HCP were conducted on a 3T Siemens Magnetom Prisma-Fit equipped with a 64-channel head coil. Structural connectivity (SC) was derived from diffusion imaging and tractography, using diffusion MRI scans acquired with a spin-echo planar imaging sequence (repetition time [TR] = 3500 ms, echo time [TE] = 64.4 ms, flip angle = 90°, refocusing flip angle = 180°, 1.6 mm isotropic voxel resolution, b-values = 300, 700, 2000 s/mm2, with 10, 40 and 90 diffusion weighing directions per shell, respectively). Functional Connectivity (FC) was derived from resting-state functional MRI (rs-fMRI), acquired with multiband accelerated 2D-BOLD echo-planar imaging (repetition [TR] = 600 ms, echo time [TE] = 30 ms, flip angle = 52°, 3 mm isotropic voxel resolution, and multiband factor of 6). For each participant, one 7 minute long scan was taken with a total of 695 frames per node. As in HCP, participants were asked to keep their eyes open. Both SC and FC data were parcellated into 200 cortical regions (nodes) using the Schaefer atlas [42].

The preprocessing of the MICA dataset was performed through the Micapipe [56]. In brief, the preprocessing of diffusion images included normalization to the mean b0 image, and correction for susceptibility distortion, head motion, and eddy currents. For each subject, probabilistic tractography was run using the iFOD2 algorithm (40 million streamlines, maximum length = 250, cutoff = 0.06). In this case, the connection weights between parcellated regions are defined as the weighted streamline count.

The preprocessing of resting state data included distortion and motion correction, registration to subjects’ respective T1-weighted data, bias and intensity normalization, as well as FSL’s ICA FIX tool trained with an in-house classifier. The data was global signal regressed in addition to the other preprocessing steps described in this pipeline.

### 4.2 Group-representative matrices

For both datasets, we generated one group-averaged covariance matrix. To do so, the time series of each node were appended across participants and runs –that means 418,000 and 34,750 time points for each node for the HCP and MICA, respectively. Then, a single covariance matrix was computed for all pairs of nodes.

Therefore, we also needed one group-averaged structural connectivity matrix. For the HCP, we constructed the distance-dependent weighted (DDW) group-representative, which was introduced in [43], as a modification of the procedure presented in [57]. This method allows us to construct a structural consensus matrix that preserves the connection distance distribution of the original subject-level connectomes, while preserving their density. Additionally, the DDW group-representative preserves the distribution of connection weights.

For MICA, the subject-level matrices available in the repository are nearly fully connected. At this density, standard network descriptors employed in this study, including degree and connected components, become weakly informative. Therefore, we averaged all subjects to construct an average group-representative network and we thresholded it by connection weight until we matched the connection density of the HCP.

The mean covariance matrices for the HCP and MICA datasets are highly correlated (*R* = 0.85, *p* = 0), whereas the SC matrices are significantly less correlated (*R* = 0.54, *p* = 0).

### 4.3 Information theory

In this work, we quantified higher-order interactions using information theory. In particular, we used the Organizational Information (or O-information), introduced by Rosas and Mediano [38]. This measure is defined as

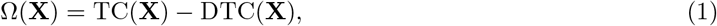

where TC(**X**) and DTC(**X**) are the total correlation [58] and dual total correlation [59], respectively. These measures are two multivariate generalizations of the mutual information, and are based on the Shannon entropy, *H*(*X*_*i*_), the fundamental information-theoretic metric that measures the amount of uncertainty inherent in the state of a variable [60].

Particularly, the Total Correlation (TC) [58] is defined as

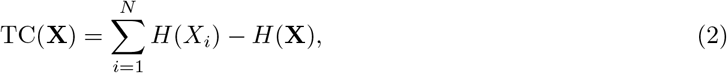

where *H*(*X*_*i*_) is the entropy of every individual variable *X*_*i*_, **X** = {*X*_1_, *X*_2_, *…, X*_*N*_} denotes a set of *N* variables, and *H*(**X**) is the joint entropy of **X**. Following this definition, TC is low when all individual variables *X*_*i*_ are independent, in which case the sum of their individual entropies equals the joint entropy. In contrast, TC is high when each variable has high entropy but the joint state of the system has low entropy, i.e. when knowing the state of one variable greatly reduces the uncertainty about the state of the others. In this way, TC increases as the system shifts from randomness toward synchrony, making it a useful measure for quantifying redundant interactions in multivariate systems.

The Dual Total Correlation (DTC) is defined as

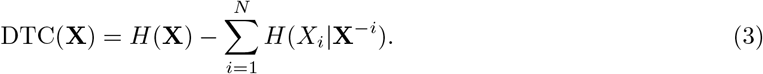

Here *H*(*X*_*i*_ | **X**^−*i*^) is the residual entropy of the variable *X*_*i*_, i.e., the uncertainty that remains about *X*_*i*_ after knowing the state of all other variables in the system. Thus, the difference in Eq. (3) quantifies the entropy that is shared by at least two elements of the system. The DTC is low when the system is either fully synchronous or completely random, and high when the variables in the system share some information in a nontrivial way.

When TC is larger than DTC, then Ω *>* 0, and the system is dominated by redundant information. Conversely, when TC is smaller than DTC, then Ω *<* 0, and synergistic dependencies predominate in the system.

Alternatively, the O-information can be expressed just in terms of total correlation as

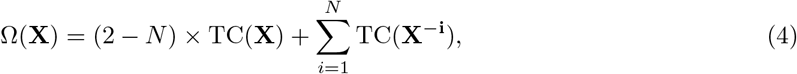

where **X**^−*i*^ is the system **X** after the *i*-th variable has been removed, see [38] for a detailed derivation. We used this formulation to calculate the O-information values of the subsets studied.

### 4.4 Information theory in Gaussian systems

It has been demonstrated that fMRI BOLD signals can be approximated by multivariate Gaussian distributions [61, 62, 63], which allows us to use the closed-form expressions of the Gaussian entropy.

For a single Gaussian variable, *X* ∼ 𝒩 (*µ, σ*^2^), the entropy is given by

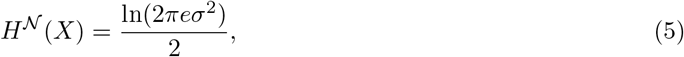

whereas when multiple variables come into consideration, **X** = *{X*_1_, *X*_2_, *…, X*_*N*_ *}*, the joint entropy is

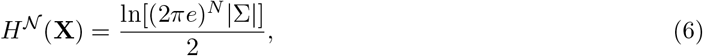

where |Σ | denotes the determinant of the covariance matrix of **X**. For a detailed derivation of these formulas see [64].

### 4.5 Consensus modules and agreement matrix

To detect network communities in the connectome, we used a multiresolution consensus clustering method [65]. The algorithm is based on the Louvain method, uses a spatial resolution parameter *γ*, and consists of three steps. First, we linearly sample the resolution parameter from two previously chosen bounds, in our case 100 samples with *γ* ∈[0, 4], and we keep the extremes of the values of *γ* that yield communities between 2 and the number of nodes in the network. Second, we perform a finer sweep, with 1000 samples, over the range of selected *γ* in the previous step. The resulting partitions are used to compute a co-classification or agreement matrix, whose elements indicate the probability with which any two nodes were assigned to the same community. Finally, we subtract from the co-classification matrix a scalar null model that estimates the expected co-classification probability under size-preserving random community assignment. The adjusted co-classification matrix is then re-clustered under a variable consensus threshold, *τ* [66], which we set to *τ* = 0.25.

## Acknowledgments

L. B. acknowledges support from the Ministry of Universities of Spain in the form of the FPU predoctoral contract as well as from the AccelNet-MultiNet program, a project of the National Science Foundation (Awards #1927425 and #1927418). M. Á. S and L. B. acknowledge support from grant PID2022-137505NB-C22 funded by MICIU/AEI/10.13039/501100011033 and by ERDF/EU. M. P. is supported by the National Science Foundation Graduate Research Fellowship under Grant No. 2240777. Any opinion, findings, and conclusions or recommendations expressed in this material are those of the authors and do not necessarily reflect the views of the National Science Foundation. This research was supported in part by Lilly Endowment, Inc., through its support for the Indiana University Pervasive Technology Institute which provided the computer infrastructure required for completion of the project. The funders had no role in study design, data collection and analysis, decision to publish, or preparation of the manuscript. Data were provided, in part, by the Human Connectome Project, WU-Minn Consortium (principal investigators: D. Van Essen and K. Ugurbil; 1U54MH091657), funded by the 16 NIH Institutes and centers that support the NIH Blueprint for Neuroscience Research and by the McDonnell Center for Systems Neuroscience at Washington University.

## References

[1] Arnaud Messé, David Rudrauf, Habib Benali, and Guillaume Marrelec. Relating structure and function in the human brain: relative contributions of anatomy, stationary dynamics, and non-stationarities. PLoS Computational Biology, 10(3):e1003530, 2014.

[2] Arnaud Messé, David Rudrauf, Alain Giron, and Guillaume Marrelec. Predicting functional connectivity from structural connectivity via computational models using mri: an extensive comparison study. NeuroImage, 111:65–75, 2015.

[3] Guillaume Marrelec, Arnaud Messé, Alain Giron, and David Rudrauf. Functional connectivity’s degenerate view of brain computation. PLoS Computational Biology, 12(10):e1005031, 2016.

[4] Laura E Súarez, Ross D Markello, Richard F Betzel, and Bratislav Misic. Linking structure and function in macroscale brain networks. Trends in Cognitive Sciences, 24(4):302–315, 2020.

[5] Gerald M. Edelman and Joseph A. Gally. Degeneracy and complexity in biological systems. Proceedings of the National Academy of Sciences, 98(24):13763–13768, 2001.

[6] Sophie Benitez Stulz, Samy Castro, Gregory Dumont, Boris Gutkin, and Demian Battaglia. Compensating functional connectivity changes due to structural connectivity damage via modifications of local dynamics. bioRxiv, 2024.

[7] A Ghosh, Y Rho, AR McIntosh, R Kötter, and VK Jirsa. Cortical network dynamics with time delays reveals functional connectivity in the resting brain. Cognitive Neurodynamics, 2(2):115–120, 2008.

[8] Gustavo Deco, Viktor Jirsa, Olaf Sporns, and Rolf Kötter. Key role of coupling, delay, and noise in resting brain fluctuations. Proceedings of the National Academy of Sciences, 106(25):10302–10307, 2009.

[9] Christopher J Honey, Olaf Sporns, Leila Cammoun, Xavier Gigandet, Jean-Philippe Thiran, Reto Meuli, and Patric Hagmann. Predicting human resting-state functional connectivity from structural connectivity. Proceedings of the National Academy of Sciences, 106(6):2035–2040, 2009.

[10] Gustavo Deco and Viktor K Jirsa. Ongoing cortical activity at rest: criticality, multistability, and ghost attractors. Journal of Neuroscience, 32(10):3366–3375, 2012.

[11] Joana Cabral, Morten L Kringelbach, and Gustavo Deco. Exploring the network dynamics underlying brain activity during rest. Progress in Neurobiology, 114:102–131, 2014.

[12] Joana Cabral, Morten L Kringelbach, and Gustavo Deco. Functional connectivity dynamically evolves on multiple time-scales over a static structural connectome: Models and mechanisms. NeuroImage, 160:84–96, 2017.

[13] Makoto Fukushima and Olaf Sporns. Comparison of fluctuations in global network topology of modeled and empirical brain functional connectivity. PLoS Computational Biology, 14(9):e1006497, 2018.

[14] Makoto Fukushima and Olaf Sporns. Structural determinants of dynamic fluctuations between segregation and integration on the human connectome. Communications Biology, 3(1):606, 2020.

[15] Maria Pope, Makoto Fukushima, Richard F Betzel, and Olaf Sporns. Modular origins of high-amplitude cofluctuations in fine-scale functional connectivity dynamics. Proceedings of the National Academy of Sciences, 118(46):e2109380118, 2021.

[16] Caio Seguin, Olaf Sporns, and Andrew Zalesky. Brain network communication: concepts, models and applications. Nature Reviews Neuroscience, 24(9):557–574, 2023.

[17] Maria Pope, Caio Seguin, Thomas F Varley, Joshua Faskowitz, and Olaf Sporns. Co-evolving dynamics and topology in a coupled oscillator model of resting brain function. NeuroImage, 277:120266, 2023.

[18] Kelly Shen, Gleb Bezgin, R Matthew Hutchison, Joseph S Gati, Ravi S Menon, Stefan Everling, and Anthony R McIntosh. Information processing architecture of functionally defined clusters in the macaque cortex. Journal of Neuroscience, 32(48):17465–17476, 2012.

[19] Martijn P Van Den Heuvel, Rene CW Mandl, Rene S Kahn, and Hilleke E Hulshoff Pol. Functionally linked resting-state networks reflect the underlying structural connectivity architecture of the human brain. Human Brain Mapping, 30(10):3127–3141, 2009.

[20] Giovanni Petri, Paul Expert, Federico Turkheimer, Robin Carhart-Harris, David Nutt, Peter J Hellyer, and Francesco Vaccarino. Homological scaffolds of brain functional networks. Journal of The Royal Society Interface, 11(101):20140873, 2014.

[21] Chad Giusti, Robert Ghrist, and Danielle S Bassett. Two’s company, three (or more) is a simplex: Algebraic-topological tools for understanding higher-order structure in neural data. Journal of Computational Neuroscience, 41(1):1–14, 2016.

[22] Manish Saggar, Olaf Sporns, Javier Gonzalez-Castillo, Peter A Bandettini, Gunnar Carlsson, Gary Glover, and Allan L Reiss. Towards a new approach to reveal dynamical organization of the brain using topological data analysis. Nature Communications, 9(1):1399, 2018.

[23] Andrea Santoro, Federico Battiston, Giovanni Petri, and Enrico Amico. Higher-order organization of multivariate time series. Nature Physics, 19(2):221–229, 2023.

[24] Mengsen Zhang, Samir Chowdhury, and Manish Saggar. Temporal mapper: Transition networks in simulated and real neural dynamics. Network Neuroscience, 7(2):431–460, 2023.

[25] Nicholas Timme, Wesley Alford, Benjamin Flecker, and John M Beggs. Synergy, redundancy, and multivariate information measures: an experimentalist’s perspective. Journal of Computational Neuroscience, 36(2):119–140, 2014.

[26] Marilyn Gatica, Rodrigo Cofré, Pedro AM Mediano, Fernando E Rosas, Patricio Orio, Ibai Diez, Stephan P Swinnen, and Jesus M Cortes. High-order interdependencies in the aging brain. Brain Connectivity, 11(9):734–744, 2021.

[27] Marilyn Gatica, Fernando E. Rosas, Pedro AM Mediano, Ibai Diez, Stephan P. Swinnen, Patricio Orio, Rodrigo Cofré, and Jesus M. Cortes. High-order functional redundancy in ageing explained via alterations in the connectome in a whole-brain model. PLoS Computational Biology, 18(9):e1010431, 2022.

[28] Andrea I Luppi, Pedro AM Mediano, Fernando E Rosas, Negin Holland, Tim D Fryer, John T O’Brien, James B Rowe, David K Menon, Daniel Bor, and Emmanuel A Stamatakis. A synergistic core for human brain evolution and cognition. Nature Neuroscience, 25(6):771–782, 2022.

[29] Simone Valenti, Laura Sparacino, Riccardo Pernice, Daniele Marinazzo, Hannes Almgren, Albert Comelli, and Luca Faes. Assessing high-order interdependencies through static o-information measures computed on resting state fmri intrinsic component networks. In International Conference on Image Analysis and Processing, pages 386–397. Springer, 2022.

[30] Thomas F Varley, Maria Pope, Joshua Faskowitz, and Olaf Sporns. Multivariate information theory uncovers synergistic subsystems of the human cerebral cortex. Communications Biology, 6(1):451, 2023.

[31] Thomas F Varley, Olaf Sporns, Stefan Schaffelhofer, Hansjörg Scherberger, and Benjamin Dann. Information-processing dynamics in neural networks of macaque cerebral cortex reflect cognitive state and behavior. Proceedings of the National Academy of Sciences, 120(2):e2207677120, 2023.

[32] Loren Koçillari, Marco Celotto, Nikolas A Francis, Shoutik Mukherjee, Behtash Babadi, Patrick O Kanold, and Stefano Panzeri. Behavioural relevance of redundant and synergistic stimulus information between functionally connected neurons in mouse auditory cortex. Brain Informatics, 10(1):34, 2023.

[33] Marilyn Gatica, Cyril Atkinson-Clement, Pedro AM Mediano, Mohammad Alkhawashki, James Ross, Jérôme Sallet, and Marcus Kaiser. Transcranial ultrasound stimulation effect in the redundant and synergistic networks consistent across macaques. Network Neuroscience, 8(4):1032–1050, 2024.

[34] Maria Pope, Thomas F Varley, Maria Grazia Puxeddu, Joshua Faskowitz, and Olaf Sporns. Time-varying synergy/redundancy dominance in the human cerebral cortex. Journal of Physics: Complexity, 6(1):015015, 2025.

[35] Thomas F Varley, Olaf Sporns, Nathan J Stevenson, Pauliina Yrjölä, Martha G Welch, Michael M Myers, Sampsa Vanhatalo, and Anton Tokariev. Emergence of a synergistic scaffold in the brains of human infants. Communications Biology, 8(1):743, 2025.

[36] Maria Grazia Puxeddu, Maria Pope, Thomas F Varley, Joshua Faskowitz, and Olaf Sporns. Leveraging multivariate information for community detection in functional brain networks. Communications Biology, 8(1):840, 2025.

[37] Samantha Pescevich Sherrill. From Spikes to Synergy: Features of Neural Computation in Local Cortical Networks. Indiana University, 2021.

[38] Fernando E Rosas, Pedro AM Mediano, Michael Gastpar, and Henrik J Jensen. Quantifying high-order interdependencies via multivariate extensions of the mutual information. Physical Review E, 100(3):032305, 2019.

[39] Borja Camino-Pontes, Antonio Jimenez-Marin, Iñigo Tellaetxe-Elorriaga, Izaro Fernandez-Iriondo, Asier Erramuzpe, Ibai Diez, Paolo Bonifazi, Marilyn Gatica, Fernando E Rosas, Daniele Marinazzo, et al. Brain structural modules associated to functional high-order interactions in the human brain. bioRxiv, pages 2025–03, 2025.

[40] David C Van Essen, Stephen M Smith, Deanna M Barch, Timothy EJ Behrens, Essa Yacoub, Kamil Ugurbil, Wu-Minn HCP Consortium, et al. The wu-minn human connectome project: an overview. NeuroImage, 80:62–79, 2013.

[41] Jessica Royer, Raúl Rodríguez-Cruces, Shahin Tavakol, Sara Larivière, Peer Herholz, Qiongling Li, Reinder Vos de Wael, Casey Paquola, Oualid Benkarim, Bo-yong Park, et al. An open mri dataset for multiscale neuroscience. Scientific Data, 9(1):569, 2022.

[42] Alexander Schaefer, Ru Kong, Evan M Gordon, Timothy O Laumann, Xi-Nian Zuo, Avram J Holmes, Simon B Eickhoff, and BT Thomas Yeo. Local-global parcellation of the human cerebral cortex from intrinsic functional connectivity mri. Cerebral cortex, 28(9):3095–3114, 2018.

[43] Laia Barjuan, Jordi Soriano, and M Ángeles Serrano. Optimal navigability of weighted human brain connectomes in physical space. NeuroImage, 297:120703, 2024.

[44] Nicholas Metropolis, Arianna W. Rosenbluth, Marshall N. Rosenbluth, Augusta H. Teller, and Edward Teller. Equation of state calculations by fast computing machines. The Journal of Chemical Physics, 21:1087–1092, 1953.

[45] Pedro AM Mediano, Fernando E Rosas, Andrea I Luppi, Robin L Carhart-Harris, Daniel Bor, Anil K Seth, and Adam B Barrett. Toward a unified taxonomy of information dynamics via integrated information decomposition. Proceedings of the National Academy of Sciences, 122(39):e2423297122, 2025.

[46] Thomas Robiglio, Matteo Neri, Davide Coppes, Cosimo Agostinelli, Federico Battiston, Maxime Lucas, and Giovanni Petri. Synergistic signatures of group mechanisms in higher-order systems. Physical Review Letters, 134(13):137401, 2025.

[47] Fernando E Rosas, Pedro AM Mediano, Andrea I Luppi, Thomas F Varley, Joseph T Lizier, Sebastiano Stramaglia, Henrik J Jensen, and Daniele Marinazzo. Disentangling high-order mechanisms and high-order behaviours in complex systems. Nature Physics, 18(5):476–477, 2022.

[48] Paul L Williams and Randall D Beer. Nonnegative decomposition of multivariate information. arXiv preprint arXiv:1004.2515, 2010.

[49] Robin AA Ince. The partial entropy decomposition: Decomposing multivariate entropy and mutual information via pointwise common surprisal. arXiv preprint arXiv:1702.01591, 2017.

[50] Thomas F Varley. Generalized decomposition of multivariate information. PLOS One, 19(2):e0297128, 2024.

[51] Matthew F Glasser, Stamatios N Sotiropoulos, J Anthony Wilson, Timothy S Coalson, Bruce Fischl, Jesper L Andersson, Junqian Xu, Saad Jbabdi, Matthew Webster, Jonathan R Polimeni, et al. The minimal preprocessing pipelines for the human connectome project. NeuroImage, 80:105–124, 2013.

[52] Douglas N Greve and Bruce Fischl. Accurate and robust brain image alignment using boundary-based registration. NeuroImage, 48(1):63–72, 2009.

[53] Hiromasa Takemura, Cesar F Caiafa, Brian A Wandell, and Franco Pestilli. Ensemble tractography. PLoS Computational Biology, 12(2):e1004692, 2016.

[54] Emma C Robinson, Saad Jbabdi, Matthew F Glasser, Jesper Andersson, Gregory C Burgess, Michael P Harms, Stephen M Smith, David C Van Essen, and Mark Jenkinson. Msm: a new flexible framework for multimodal surface matching. NeuroImage, 100:414–426, 2014.

[55] Linden Parkes, Ben Fulcher, Murat Yücel, and Alex Fornito. An evaluation of the efficacy, reliability, and sensitivity of motion correction strategies for resting-state functional mri. NeuroImage, 171:415–436, 2018.

[56] Raúl R Cruces, Jessica Royer, Peer Herholz, Sara Larivière, Reinder Vos De Wael, Casey Paquola, Oualid Benkarim, Bo-yong Park, Janie Degré-Pelletier, Mark C Nelson, et al. Micapipe: A pipeline for multimodal neuroimaging and connectome analysis. NeuroImage, 263:119612, 2022.

[57] Richard F Betzel, Alessandra Griffa, Patric Hagmann, and Bratislav Mišić. Distance-dependent consensus thresholds for generating group-representative structural brain networks. Network Neuroscience, 3(2):475–496, 2019.

[58] Satosi Watanabe. Information theoretical analysis of multivariate correlation. IBM Journal of Research and Development, 4(1):66–82, 1960.

[59] Te Sun Han. Linear dependence structure of the entropy space. Inf. Control, 29(4):337–368, 1975.

[60] Claude Elwood Shannon. A mathematical theory of communication. The Bell System Technical Journal, 27:379–423, 1948.

[61] Jaroslav Hlinka, Milan Paluš, Martin Vejmelka, Dante Mantini, and Maurizio Corbetta. Functional connectivity in resting-state fmri: is linear correlation sufficient? NeuroImage, 54(3):2218–2225, 2011.

[62] Raphaël Liégeois, BT Thomas Yeo, and Dimitri Van De Ville. Interpreting null models of resting-state functional mri dynamics: not throwing the model out with the hypothesis. NeuroImage, 243:118518, 2021.

[63] Marc-Andre Schulz, BT Thomas Yeo, Joshua T Vogelstein, Janaina Mourao-Miranada, Jakob N Kather, Konrad Kording, Blake Richards, and Danilo Bzdok. Different scaling of linear models and deep learning in ukbiobank brain images versus machine-learning datasets. Nature Communications, 11(1):4238, 2020.

[64] Thomas M. Cover and Joy A. Thomas. Elements of Information Theory. John Wiley & Sons, New York, 1991. Online ISBN: 0-471-20061-1.

[65] Lucas GS Jeub, Olaf Sporns, and Santo Fortunato. Multiresolution consensus clustering in networks. Scientific reports, 8(1):3259, 2018.

[66] Andrea Lancichinetti and Santo Fortunato. Consensus clustering in complex networks. Scientific reports, 2(1):336, 2012.

