## Supplementary Information for "Structural signatures of synergy and redundancy in human brain function"

#### 1 HCP Dataset

| Subset size | Synergistic | Redundant | Random |
|---|---|---|---|
| 3 | 199 | 249 | 4994 |
| 4 | 577 | 273 | 5000 |
| 5 | 995 | 242 | 5000 |
| 6 | 1582 | 185 | 5000 |
| 7 | 2438 | 146 | 5000 |
| 8 | 3146 | 93 | 5000 |
| 9 | 3663 | 89 | 5000 |
| 10 | 4021 | 95 | 5000 |
| 11 | 4350 | 80 | 5000 |
| 12 | 4596 | 91 | 5000 |
| 13 | 4759 | 84 | 5000 |
| 14 | 4843 | 89 | 5000 |
| 15 | 4889 | 99 | 5000 |

Table ST1: Number of unique subsets found by the optimization algorithm for each subset size and subset type, for the HCP dataset.

| | **Comparison** | **Size 3** | | **Size 6** | | **Size 10** | |
|---|---|---|---|---|---|---|---|
| | | **D** | **p-value** | **D** | **p-value** | **D** | **p-value** |
| Number components | SYN vs RED | 0.22 | 0 | 0.47 | 0 | 0.48 | 0 |
| | SYN vs RAND | -0.70 | 0 | -0.77 | 0 | -0.29 | 0 |
| | RED vs RAND | -0.92 | 0 | -1.24 | 0 | -0.76 | 0 |
| Connection density | SYN vs RED | 0.20 | 0 | 0.48 | 0 | 0.83 | 0 |
| | SYN vs RAND | 0.39 | 0 | 0.41 | 0 | 0.14 | 0 |
| | RED vs RAND | 0.58 | 0 | 0.78 | 0 | 0.84 | 0 |
| Connection weight | SYN vs RED | 0.30 | 0 | 0.58 | 0 | 0.59 | 0 |
| | SYN vs RAND | 0.43 | 0 | 0.35 | 0 | 0.47 | 0 |
| | RED vs RAND | 0.54 | 0 | 0.70 | 0 | 0.78 | 0 |

Table ST2: Summary of distributional statistics in Section *Subset on its own.* We report the statistic D, defined as the KS statistic for continuous distributions and as the difference in means for discrete distributions, together with the corresponding p-values.

| | Comparison | Size 3 | | Size 6 | | Size 10 | |
|---|---|---|---|---|---|---|---|
| | | **D** | **p-value** | **D** | **p-value** | **D** | **p-value** |
| Degree | SYN vs RED | 0.55 | 0 | 0.59 | 0 | 0.40 | 0 |
| | SYN vs RAND | 0.42 | 0 | 0.37 | 0 | 0.15 | 0 |
| | RED vs RAND | 0.32 | 0 | 0.40 | 0 | 0.30 | 0 |
| Strength | SYN vs RED | 0.19 | 7e-04 | 0.16 | 1e-04 | 0.40 | 0 |
| | SYN vs RAND | 0.33 | 0 | 0.26 | 0 | 0.24 | 0 |
| | RED vs RAND | 0.16 | 0 | 0.26 | 0 | 0.47 | 0 |
| Betweenness centrality | SYN vs RED | 0.29 | 0 | 0.37 | 0 | 0.41 | 0 |
| | SYN vs RAND | 0.23 | 0 | 0.22 | 0 | 0.14 | 0 |
| | RED vs RAND | 0.13 | 7e-04 | 0.15 | 2e-04 | 0.29 | 0 |
| Clustering | SYN vs RED | 0.44 | 0 | 0.54 | 0 | 0.38 | 0 |
| | SYN vs RAND | 0.23 | 0 | 0.24 | 0 | 0.14 | 0 |
| | RED vs RAND | 0.30 | 0 | 0.37 | 0 | 0.28 | 0 |
| Number communities | SYN vs RED | 0.36 | 0 | 1.41 | 0 | 2.65 | 0 |
| | SYN vs RAND | -0.33 | 0 | -0.54 | 0 | -0.42 | 0 |
| | RED vs RAND | -0.69 | 0 | -1.94 | 0 | -3.06 | 0 |
| Stability assignment | SYN vs RED | 0.58 | 0 | 0.62 | 0 | 0.41 | 0 |
| | SYN vs RAND | 0.31 | 0 | 0.30 | 0 | 0.08 | 0 |
| | RED vs RAND | 0.33 | 0 | 0.41 | 0 | 0.34 | 0 |

Table ST3: Summary of distributional statistics in Section *Subset as part of the network*. We report the statistic D, defined as the KS statistic for continuous distributions and as the difference in means for discrete distributions, together with the corresponding p-values.

| | Comparison | Size 3 | | Size 6 | | Size 10 | |
|---|---|---|---|---|---|---|---|
| | | **D** | **p-value** | **D** | **p-value** | **D** | **p-value** |
| Connection density | SYN vs RED | 0.56 | 0 | 0.63 | 0 | 0.51 | 0 |
| | SYN vs RAND | 0.40 | 0 | 0.36 | 0 | 0.14 | 0 |
| | RED vs RAND | 0.34 | 0 | 0.47 | 0 | 0.42 | 0 |
| Fraction nodes reached | SYN vs RED | 0.53 | 0 | 0.70 | 0 | 0.78 | 0 |
| | SYN vs RAND | 0.14 | 4e-04 | 0.18 | 0 | 0.12 | 0 |
| | RED vs RAND | 0.59 | 0 | 0.77 | 0 | 0.79 | 0 |
| Jaccard index | SYN vs RED | 0.11 | 1.219e-01 | 0.36 | 0 | 0.86 | 0 |
| | SYN vs RAND | 0.53 | 0 | 0.52 | 0 | 0.29 | 0 |
| | RED vs RAND | 0.59 | 0 | 0.77 | 0 | 0.89 | 0 |

Table ST4: Summary of distributional statistics in Section *Interface between subset and rest of the network*. We report the statistic D, defined as the KS statistic for continuous distributions and as the difference in means for discrete distributions, together with the corresponding p-values.

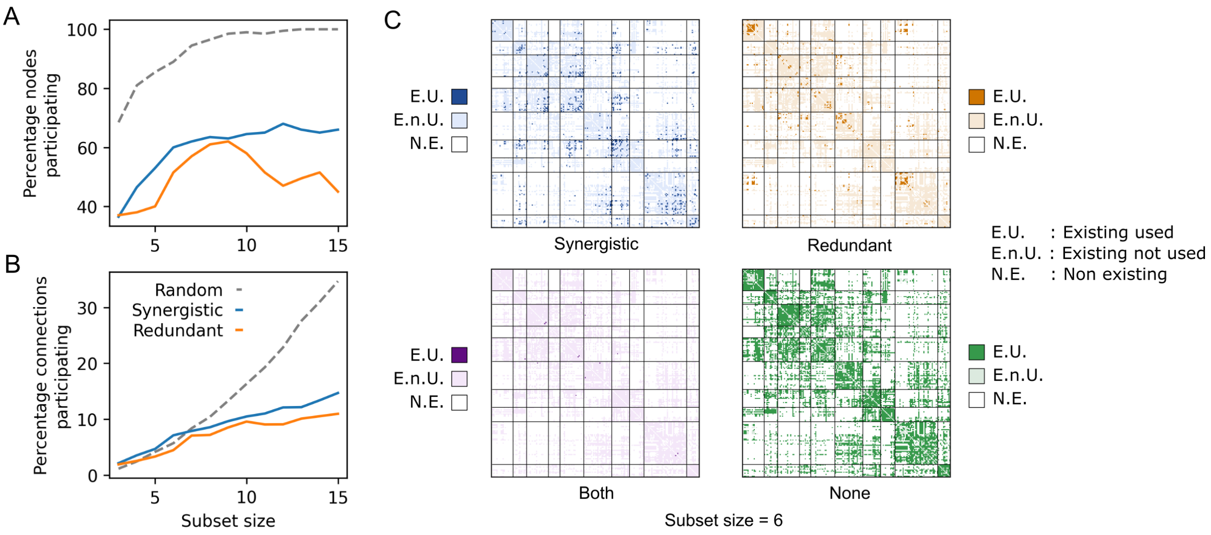


Figure SF1: **Participation of nodes and connections in optimized subsets only considering 80 subsets.** Figure calculated with only 80 subsets for each subset type. Percentage of A) nodes and B) connections participating at least once in each type of subset across all subset sizes. Results for synergistic, redundant and random subsets are shown by the solid blue line, solid orange line and dashed gray line, respectively. C) Instances of the SC matrix where connections that meet a certain criteria are highlighted: in the top-left panel, connections participating in synergistic subsets; in the top-right, those in redundant subsets; in the bottom-left, those in both; and in the bottom-right, those that never participate. In all panels, non-existent connections are shown in white, existing but non-highlighted connections in light colors, and highlighted connections in bright colors. Results correspond to subsets with 6 nodes.

### 2 MICA Dataset

| Subset size | Synergistic | Redundant | Random |
|---|---|---|---|
| 3 | 243 | 241 | 4994 |
| 4 | 560 | 190 | 5000 |
| 5 | 859 | 91 | 5000 |
| 6 | 1368 | 67 | 5000 |
| 7 | 2171 | 70 | 5000 |
| 8 | 2973 | 53 | 5000 |
| 9 | 3624 | 59 | 5000 |
| 10 | 4166 | 53 | 5000 |
| 11 | 4503 | 42 | 5000 |
| 12 | 4716 | 37 | 5000 |
| 13 | 4863 | 44 | 5000 |
| 14 | 4902 | 53 | 5000 |
| 15 | 4948 | 54 | 5000 |

Table ST5: Number of unique subsets found by the optimization algorithm for each subset size and subset type, for the MICA dataset.

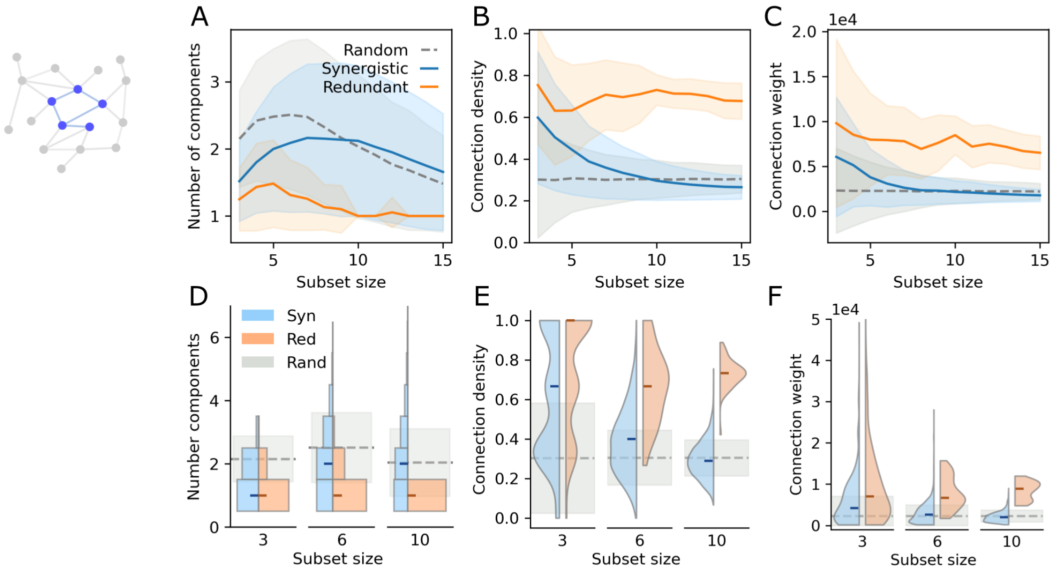


Figure SF2: **Subset on its own.** A-C) Mean results of number of components within the subgraph A), connection density B), and average connection weight C), across all subset sizes. The results for synergistic subsets are represented by the solid blue line; for redundant subsets by the solid orange line; and for random subsets by the dashed gray line. The shadowed areas correspond to the standard deviation in each case. D-F) Distributions of values of number of components D), connection density E) and average connection weight F), for subsets with 3, 6 and 10 nodes. Values for synergistic and redundant subsets are depicted in blue and orange, respectively. The median value of each distribution is indicated by a small horizontal mark. The mean and standard deviation for random subsets are represented by the dashed gray line and the shadowed gray area, respectively.

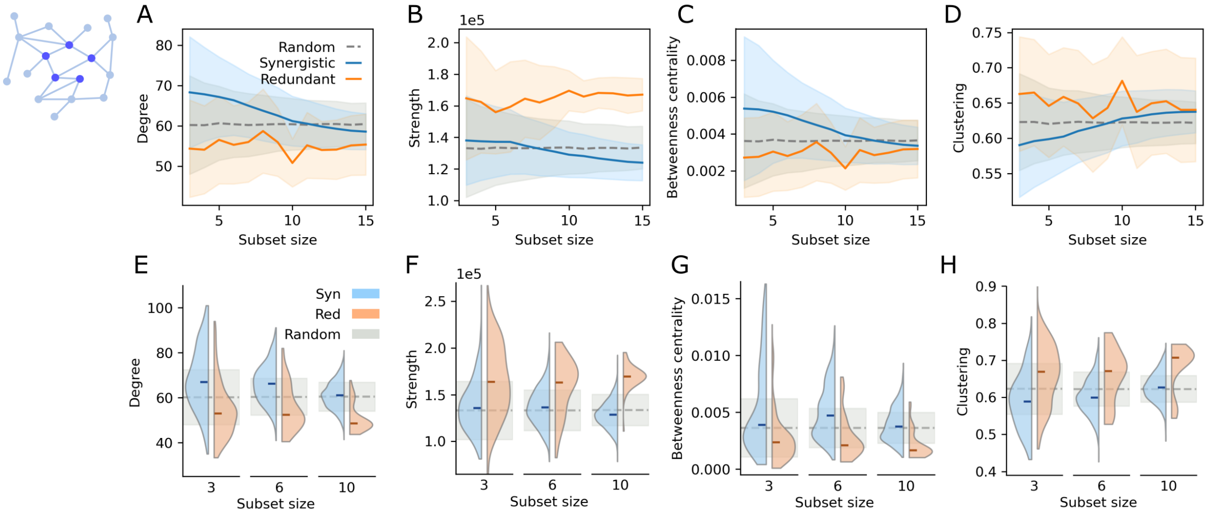


Figure SF3: **Subset as part of the network.** A-D) Mean results of average A) degree, B) strength, C) betweenness centrality and D) clustering coefficient, across all subset sizes. The results for synergistic subsets are represented by the solid blue line; for redundant subsets by the solid orange line; and for random subsets by the dashed gray line. The shadowed areas correspond to the standard deviation in each case. E-H) Distributions of average E) degree, F) strength, G) betweenness centrality and H) clustering coefficient, for subsets with 3, 6 and 10 nodes. Values for synergistic and redundant subsets, depicted in blue and orange, respectively. The median value of each distribution is indicated by a small horizontal mark. The mean and standard deviation for nodes in random subsets are represented by the dashed gray line and the shadowed gray area, respectively.

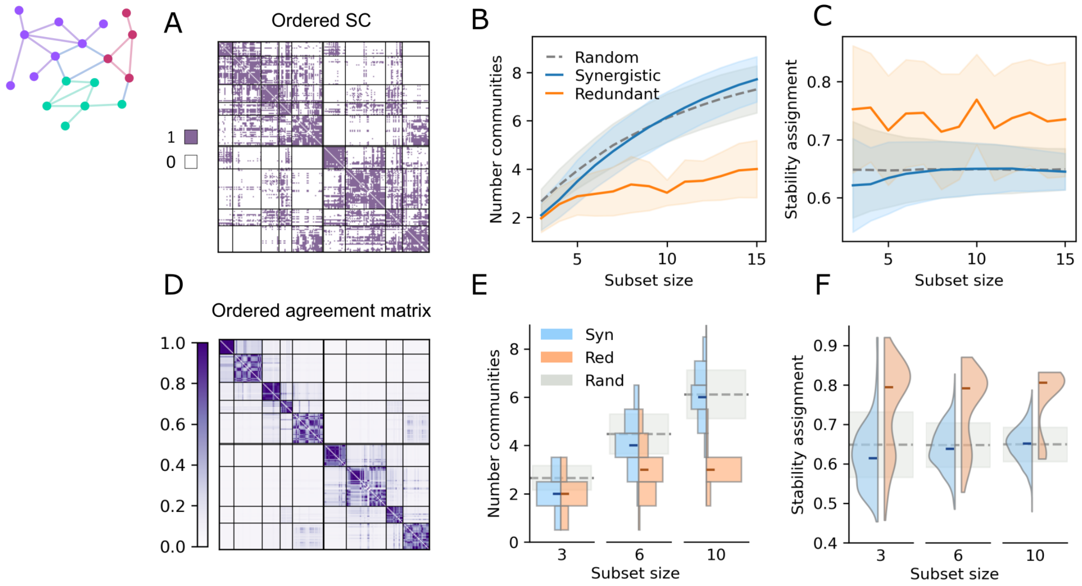


Figure SF4: **Community structure.** A) Structural connectivity matrix ordered by structural communities, whose boundaries are indicated by black solid lines. Existing connections are shown in purple; non-existing connections in white. B-C) Mean results of number of communities appearing in each type of subset and of node community assignment stability, respectively. The results for synergistic subsets are represented by the solid blue line; for redundant subsets by the solid orange line; and for random subsets by the dashed gray line. The shadowed areas correspond to the standard deviation in each case. D) Agreement matrix ordered by structural communities, whose boundaries are indicated by black solid lines. Darker colors represent higher agreement. For subsets with 3, 6 and 10 nodes: distributions of E) number of communities appearing in each type of subset and F) average node community assignment stability. Results for synergy and redundancy are shown in blue and orange, respectively. The mean and standard deviation for random subsets are represented by the dashed gray line and the shadowed gray area, respectively.

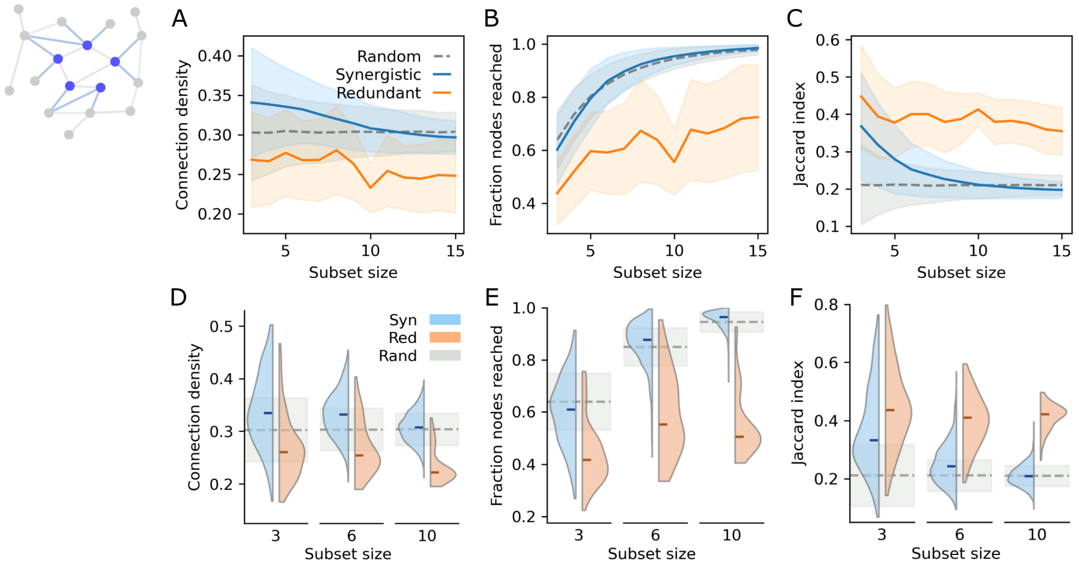


Figure SF5: **Interface between subset and rest of network.** A-C) Mean results of A) subset-to-network connection density in the interface, B) fraction of nodes of the network that are one connection away from the nodes in the subset, and C) average Jaccard index between neighbor patterns. The results for synergistic subsets are represented by the solid blue line; for redundant subsets by the solid orange line; and for random subsets by the dashed gray line. The shadowed areas correspond to the standard deviation in each case. For subsets with 3, 6 and 10 nodes: distributions of D) subset-to-network connection density in the interface, E) fraction of nodes of the network that are one connection away from the nodes in the subset, and F) average Jaccard index between neighbor patterns. Results for synergistic and redundant subsets are shown in blue and orange, respectively. The median value of each distribution is indicated by a small horizontal mark. For random subsets, the mean is indicated by the dashed gray line and the standard deviation by the shadowed gray area.

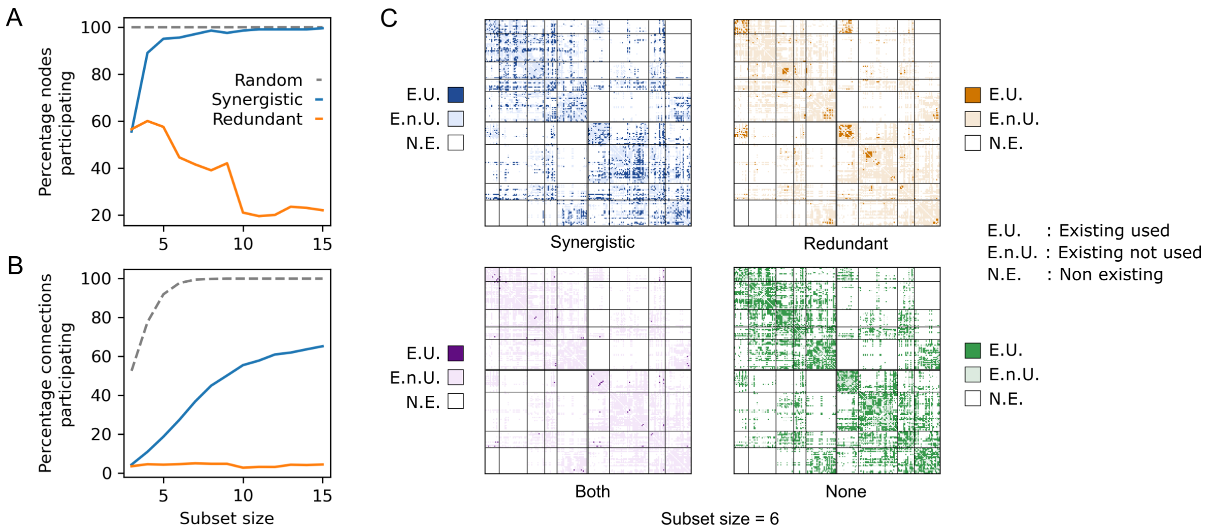


Figure SF6: **Participation of nodes and connections in optimized subsets.** Percentage of A) nodes and B) connections participating at least once in each type of subset across all subset sizes. Results for synergistic, redundant and random subsets are shown by the solid blue line, solid orange line and dashed gray line, respectively. C) Instances of the SC matrix where connections that meet a certain criteria are highlighted: in the top-left panel, connections participating in synergistic subsets; in the top-right, those in redundant subsets; in the bottom-left, those in both; and in the bottom-right, those that never participate. In all panels, non-existent connections are shown in white, existing but non-highlighted connections in light colors, and highlighted connections in bright colors. Results correspond to subsets with 6 nodes.

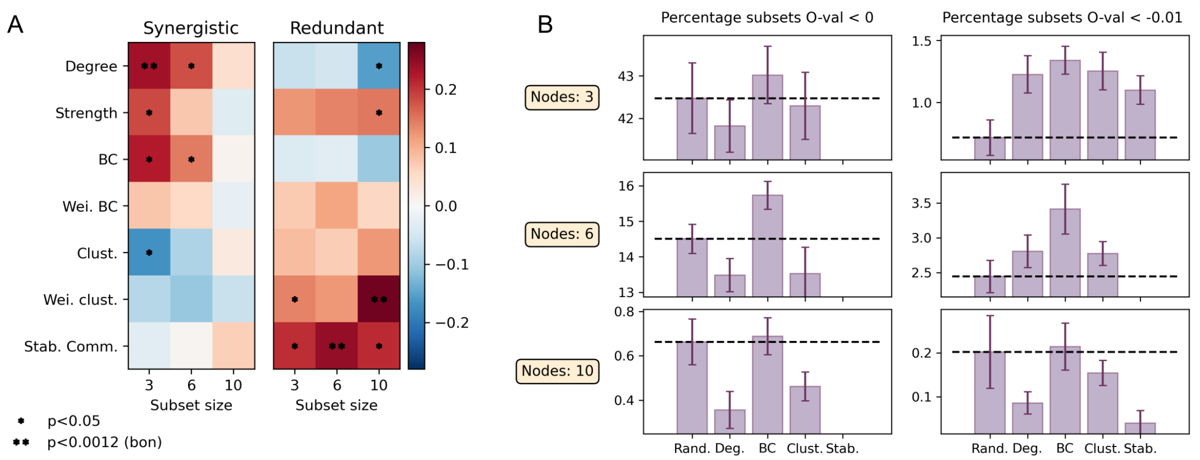


Figure SF7: **Node participation frequency.** A) For synergistic and redundant subsets with 3, 6 and 10 nodes: Spearman correlation between node participation frequency and degree, strength, betweenness centrality (BC), weighted betweenness centrality, clustering, weighted clustering and stability in the community assignment. Positive and negative correlations are shown in red and blue, respectively. Entries with p-values smaller than 0.05 are marked with an asterisk. Entries with p-values smaller than the Bonferroni-corrected threshold are marked with two asterisks. B) Percentage of synergistic subsets obtained biasing the selection process by degree (Deg.), betweenness centrality (BC), clustering (Clust.) and stability in the community assignment (Stab.) with respect to random selection. The limits of the y-axis have been adjusted to visualize better the differences between bars.

| | **Comparison** | **Size 3** | | **Size 6** | | **Size 10** | |
|---|---|---|---|---|---|---|---|
| | | **D** | **p-value** | **D** | **p-value** | **D** | **p-value** |
| Number components | SYN vs RED | 0.27 | 0 | 0.78 | 0 | 1.12 | 0 |
| | SYN vs RAND | -0.63 | 0 | -0.42 | 0 | 0.08 | 6e-04 |
| | RED vs RAND | -0.90 | 0 | -1.19 | 0 | -1.04 | 0 |
| Connection density | SYN vs RED | 0.23 | 0 | 0.56 | 0 | 0.98 | 0 |
| | SYN vs RAND | 0.34 | 0 | 0.21 | 0 | 0.04 | 4e-04 |
| | RED vs RAND | 0.57 | 0 | 0.72 | 0 | 0.98 | 0 |
| Connection weight | SYN vs RED | 0.23 | 0 | 0.60 | 0 | 0.96 | 0 |
| | SYN vs RAND | 0.44 | 0 | 0.23 | 0 | 0.06 | 0 |
| | RED vs RAND | 0.60 | 0 | 0.70 | 0 | 0.94 | 0 |

Table ST6: Summary of distributional statistics in Section *Subset on its own*. We report the statistic D, defined as the KS statistic for continuous distributions and as the difference in means for discrete distributions, together with the corresponding p-values.

| | Comparison | Size 3 | | Size 6 | | Size 10 | |
|---|---|---|---|---|---|---|---|
| | | **D** | **p-value** | **D** | **p-value** | **D** | **p-value** |
| Degree | SYN vs RED | 0.48 | 0 | 0.52 | 0 | 0.70 | 0 |
| | SYN vs RAND | 0.25 | 0 | 0.27 | 0 | 0.06 | 0 |
| | RED vs RAND | 0.27 | 0 | 0.37 | 0 | 0.68 | 0 |
| Strength | SYN vs RED | 0.35 | 0 | 0.48 | 0 | 0.90 | 0 |
| | SYN vs RAND | 0.10 | 1e-02 | 0.09 | 0 | 0.13 | 0 |
| | RED vs RAND | 0.36 | 0 | 0.51 | 0 | 0.84 | 0 |
| Betweenness centrality | SYN vs RED | 0.38 | 0 | 0.53 | 0 | 0.66 | 0 |
| | SYN vs RAND | 0.21 | 0 | 0.29 | 0 | 0.12 | 0 |
| | RED vs RAND | 0.23 | 0 | 0.38 | 0 | 0.66 | 0 |
| Clustering | SYN vs RED | 0.40 | 0 | 0.47 | 0 | 0.70 | 0 |
| | SYN vs RAND | 0.20 | 0 | 0.20 | 0 | 0.07 | 0 |
| | RED vs RAND | 0.27 | 0 | 0.40 | 0 | 0.73 | 0 |
| Number communities | SYN vs RED | 0.13 | 2e-02 | 1.19 | 0 | 3.14 | 0 |
| | SYN vs RAND | -0.57 | 0 | -0.32 | 0 | 0.04 | 4e-02 |
| | RED vs RAND | -0.70 | 0 | -1.51 | 0 | -3.10 | 0 |
| Stability assignment | SYN vs RED | 0.60 | 0 | 0.63 | 0 | 0.77 | 0 |
| | SYN vs RAND | 0.17 | 0 | 0.07 | 1e-04 | 0.02 | 2e-01 |
| | RED vs RAND | 0.54 | 0 | 0.61 | 0 | 0.77 | 0 |

Table ST7: Summary of distributional statistics in Section *Subset as part of the network*. We report the statistic D, defined as the KS statistic for continuous distributions and as the difference in means for discrete distributions, together with the corresponding p-values.

| | Comparison | Size 3 | | Size 6 | | Size 10 | |
|---|---|---|---|---|---|---|---|
| | | **D** | **p-value** | **D** | **p-value** | **D** | **p-value** |
| Connection density | SYN vs RED | 0.49 | 0 | 0.57 | 0 | 0.77 | 0 |
| | SYN vs RAND | 0.23 | 0 | 0.26 | 0 | 0.06 | 0 |
| | RED vs RAND | 0.31 | 0 | 0.45 | 0 | 0.76 | 0 |
| Fraction nodes reached | SYN vs RED | 0.55 | 0 | 0.75 | 0 | 0.94 | 0 |
| | SYN vs RAND | 0.15 | 1e-04 | 0.12 | 0 | 0.12 | 0 |
| | RED vs RAND | 0.69 | 0 | 0.73 | 0 | 0.94 | 0 |
| Jaccard index | SYN vs RED | 0.31 | 0 | 0.70 | 0 | 0.97 | 0 |
| | SYN vs RAND | 0.52 | 0 | 0.25 | 0 | 0.03 | 1e-02 |
| | RED vs RAND | 0.69 | 0 | 0.81 | 0 | 0.97 | 0 |

Table ST8: Summary of distributional statistics in Section *Interface between subset and rest of the network*. We report the statistic D, defined as the KS statistic for continuous distributions and as the difference in means for discrete distributions, together with the corresponding p-values.
